# Varicella-Zoster Virus ORF9 Is an Antagonist of the DNA Sensor cGAS

**DOI:** 10.1101/2020.02.11.943415

**Authors:** Jonny Hertzog, Wen Zhou, Rachel E. Rigby, Gerissa Fowler, Chiara Cursi, Lise Chauveau, Tamara Davenne, Philip J. Kranzusch, Jan Rehwinkel

## Abstract

Varicella-Zoster virus (VZV) causes chickenpox and shingles. Although infection is associated with severe morbidity in some individuals, molecular mechanisms that determine innate immune responses remain poorly defined. We found that the cGAS/STING DNA sensing pathway was required for type I interferon (IFN) induction during VZV infection and that recognition of VZV by cGAS restricted its replication. Viral gene overexpression screening identified the essential VZV tegument protein ORF9 as a cGAS antagonist. Ectopically and virally expressed ORF9 bound to endogenous cGAS. Confocal microscopy revealed co-localisation of cGAS and ORF9, which reduced the type I IFN response to transfected DNA. ORF9 and cGAS also interacted directly in a cell-free system and phase-separated together with DNA. Furthermore, ORF9 inhibited cGAMP production by cGAS. Taken together, we uncovered the importance of the cGAS/STING DNA sensing pathway for VZV recognition and identified a VZV immune antagonist that directly interferes with DNA sensing *via* cGAS.

## INTRODUCTION

Varicella-Zoster virus (VZV) is one of nine herpes viruses that infect humans (Arvin and Gilden, 2013). VZV is an alpha-herpesvirus, closely related to herpes simplex virus (HSV) 1 and 2. It has a 125 kb dsDNA genome, the smallest of the human herpesviruses. The genome includes at least 70 open reading frames (ORFs). Primary infection causes chickenpox (*Varicella*). Like all human herpes viruses, VZV establishes life-long latency in an infected host and can reactivate as shingles (*Zoster*). Shingles is a debilitating disease with significant associated morbidity. During both primary infection and reactivation, the virus can gain access to the central nervous system and cause severe complications such as encephalitis and vasculitis (Nagel and Gilden, 2014). Despite introduction of the live-attenuated chickenpox vaccine in the early 1990s, the virus remains highly prevalent worldwide (WHO, 2014).

The type I interferon (IFN) system lies at the forefront of host defence against infectious pathogens and is indispensable for successful control of viral infections (McNab et al., 2015). The expression of type I IFNs is induced following pattern recognition receptor (PRR) activation. PRRs are a heterogenous group of proteins that can respond to a diverse array of pathogen-associated molecular patterns (PAMPs) (Brubaker et al., 2015). The recognition of viral pathogens relies to a large extent on sensing of nucleic acids (Barrat et al., 2016; Hartmann, 2017). Both endosomal toll-like receptors and dedicated cytosolic sensors are potently activated by viral RNA and DNA. The DNA sensor cyclic GMP-AMP synthase (cGAS) synthesises the second messenger 2′3′-cyclic GMP-AMP (hereafter simply cGAMP), a cyclic dinucleotide, upon direct binding to dsDNA (Ablasser and Chen, 2019). Binding of cGAMP to stimulator of IFN genes (STING) results in the activation of the transcription factors IRF3 and NF-κB via the kinases TBK1 and IKKε (Hopfner and Hornung, 2020). IRF3 and NF-κB induce the expression of type I IFNs, type III IFNs, and inflammatory cytokines.

Type I IFNs, including IFNα and IFNβ, are secreted cytokines that act in an autocrine, paracrine, or systemic manner by binding to the type I IFN receptor (IFNAR) (McNab et al., 2015). Canonical IFNAR signalling results in the phosphorylation and heterodimerisation of the transcription factors STAT1 and STAT2. After recruitment of IRF9, this protein complex drives expression of hundreds of genes, termed interferon-stimulated genes (ISGs). Among others, ISGs contain genes that encode PRRs, proteins involved in type I IFN induction and signalling, negative and positive feedback regulators, restriction factors acting directly on viruses, and proteins that are involved in adaptive immune responses (Schoggins, 2019).

Despite the fact that VZV is a highly prevalent and important human pathogen, its pathogenesis is still poorly understood. The lack of suitable small animal models that recapitulate primary infection and latency establishment has hindered the molecular characterisation of its life cycle *in vivo* (Haberthur and Messaoudi, 2013). During its dissemination in the human host, the virus infects a multitude of different cells. Infection of T cells, keratinocytes, neurons, and epithelial cells is indispensable for VZV’s life cycle (Zerboni et al., 2014). In addition, immune cells including dendritic cells (DCs), monocytes, and NK cells are capable of supporting VZV replication *in vitro* and are potentially relevant for *in vivo* spread (Abendroth et al., 2001; Campbell et al., 2018; Kennedy et al., 2019; Morrow et al., 2003; Wang et al., 2005). Current evidence suggests that type I IFNs are critical for control of VZV infection. Increased IFNα levels can be detected in the serum of patients with primary VZV infection (Arvin et al., 1986). In addition, type I IFNs limit VZV replication *in vitro* (Kim et al., 2017; Ku et al., 2016; Shakya et al., 2019; Torigo et al., 2000). However, the events that govern the cell-intrinsic recognition of the virus in the various cell types it infects and induction of the antiviral cytokine response have only begun to be elucidated *in vitro*. The DNA sensor TLR9 is partly responsible for IFNα secretion after infection of plasmacytoid DCs (Yu et al., 2011). In dermal fibroblasts, STING is required for type I and type III IFN production (Kim et al., 2017). An interesting genetic link between DNA sensing *via* RNA polymerase III and infection of the central nervous system by VZV has been uncovered recently (Carter-Timofte et al., 2018). However, a comprehensive characterisation of the role of DNA sensing during VZV infection is still lacking.

In this study we tested which nucleic acid sensors induce type I IFN expression in response to VZV infection. We show that the cGAS – STING – TBK1 – IRF3 signalling axis was responsible for antiviral cytokine expression after VZV infection. We further report the generation of a VZV open reading frame (ORF) expression library and identification of a viral cGAS antagonist. The tegument protein encoded by ORF9 curtailed activation of cGAS and subsequent synthesis of cGAMP. Mechanistically, we show that ORF9 interacted with cGAS and DNA. This resulted in decreased cGAMP and IFN production. We propose a model in which cGAS activation upon VZV infection is limited immediately after viral entry through the tegument protein ORF9.

## RESULTS

### The Type I IFN Response to VZV Requires the DNA Sensor cGAS

To identify PRRs that induce type I IFNs in response to VZV infection we used the monocytic cell line THP1. We hypothesised a role of DNA sensors in VZV infection given its identity as a DNA virus and the previously shown role of STING in recognition of VZV (Kim et al., 2017). THP1 cells, unlike many other immortalised cell lines, induce type I and type III IFNs *via* cGAS in response to DNA (Sun et al., 2013; Wu et al., 2013). Furthermore, THP1 cells are amenable to genome editing and knockout (KO) lines can be used to genetically dissect the role of individual proteins involved in pattern recognition. THP1 cells are permissive for VZV infection and propagation (Nour et al., 2011) and VZV infects primary human monocytes and macrophages *in vitro* and *in vivo* (Kennedy et al., 2019; Mainka et al., 1998). In addition to wild-type (WT) THP1 cells, we tested previously described KO lines lacking STING, TBK1, MyD88, or IFNAR2. We further generated cGAS KO, MAVS KO and IRF3 KO cells using CRISPR/Cas9 technology (see Materials and Methods and Figures S1 and S2). All THP1 KO cells were validated by immunoblotting for the absence of protein and functionally by stimulation with DNA, RNA and type I IFN (Figures S1, S2). These cells contained a secreted luciferase reporter construct under control of an IRF3-responsive promotor. Upon treatment with PMA, THP1 cells adopt a macrophage-like, adherent and highly responsive phenotype. Given the difficulties of working with cell-free VZV (Caunt and Taylor-Robinson, 2009; Chen et al., 2004), we used co-culture with VZV-infected (+VZV) MeWo cells to infect PMA-treated THP1 cells; co-culture with uninfected MeWo cells served as a control (Figure S3A). MeWo cells are a melanoma cell line that is well-established for VZV propagation. After 48 hours of co-culture, cells were harvested for RT-qPCR and immunoblotting. For all experiments, uninfected MeWo cells were additionally used as target cells. MeWo cells do not induce type I IFNs in response to VZV (Figure S3B).

To investigate whether THP1 cells induce type I IFNs in response to VZV we analysed mRNA expression levels of *IFNB1* (encodes IFNβ) and *IFI44*, an ISG. In WT THP1 cells, both transcripts were robustly induced after VZV infection (Figure S3B). Similar results were obtained using MyD88 KO and MAVS KO cells. In contrast, no transcriptional upregulation of *IFNB1* or *IFI44* was observed in THP1 cells lacking cGAS, STING, TBK1, IRF3, or IFNAR2. Moreover, immunoblot analysis showed that the transcription factors STAT1 and STAT2 were only phosphorylated in WT, MyD88 KO, and MAVS KO cells (Figure S3C). No p-STAT1 and p-STAT2 signals were observed in cells lacking cGAS, STING, TBK1, IRF3, or IFNAR2. This indicated that only WT, MyD88 KO and MAVS KO cells secreted type I IFNs in response to VZV infection. Consistently, STAT1 and RIG-I, which are both encoded by ISGs, were upregulated at protein level only in the cells showing STAT1/2 activation (Figure S3C). Importantly, we could not observe phosphorylation of STAT1/2 or increased abundance of STAT1 and RIG-I in infected MeWo cells. Western blotting with antibodies against VZV-glycoprotein E (gE)/glycoprotein I (gI) and VZV ORF62 confirmed that all cell lines became infected (Figure S3C). Determination of CXCL10 (IP-10) levels in co-culture supernatants confirmed the findings of our RT-qPCR and immunoblot analyses. WT, MyD88 KO, and MAVS KO THP1 cells produced low levels of CXCL10 at baseline and these were increased after infection with VZV (Figure S3D). No CXCL10 was detected in supernatants from uninfected cells and in samples from infected cGAS KO, STING KO, TBK1 KO, and IFNAR2 KO THP1 cells; similarly, MeWo cells did not secrete CXCL10. We could detect low levels of CXCL10 in supernatants from IRF3 KO cells but there was no increase above baseline after infection. Collectively these results suggest that in THP1 cells the type I IFN response to VZV infection required the DNA sensor cGAS and the STING – TBK1 – IRF3 signalling axis. It is therefore likely that dsDNA is the PAMP recognised in VZV-infected cells.

To dissect the role of the DNA sensor cGAS in recognition of VZV further we developed a transwell-based infection system (Figure 1A). In this setup, infected MeWo cells are first seeded on the bottom side of the transwell membrane. After adherence, THP1 target cells are seeded on the opposite side of the membrane. The membrane contains 1 μm pores through which cell-cell contacts can be established and VZV can spread. Importantly, the inoculum and target cells do not mix, and a homogenous target cell population can be harvested for analysis. We anticipate that this new infection protocol (see Methods for details) will be widely applicable to many VZV research projects.

**Figure 1:**
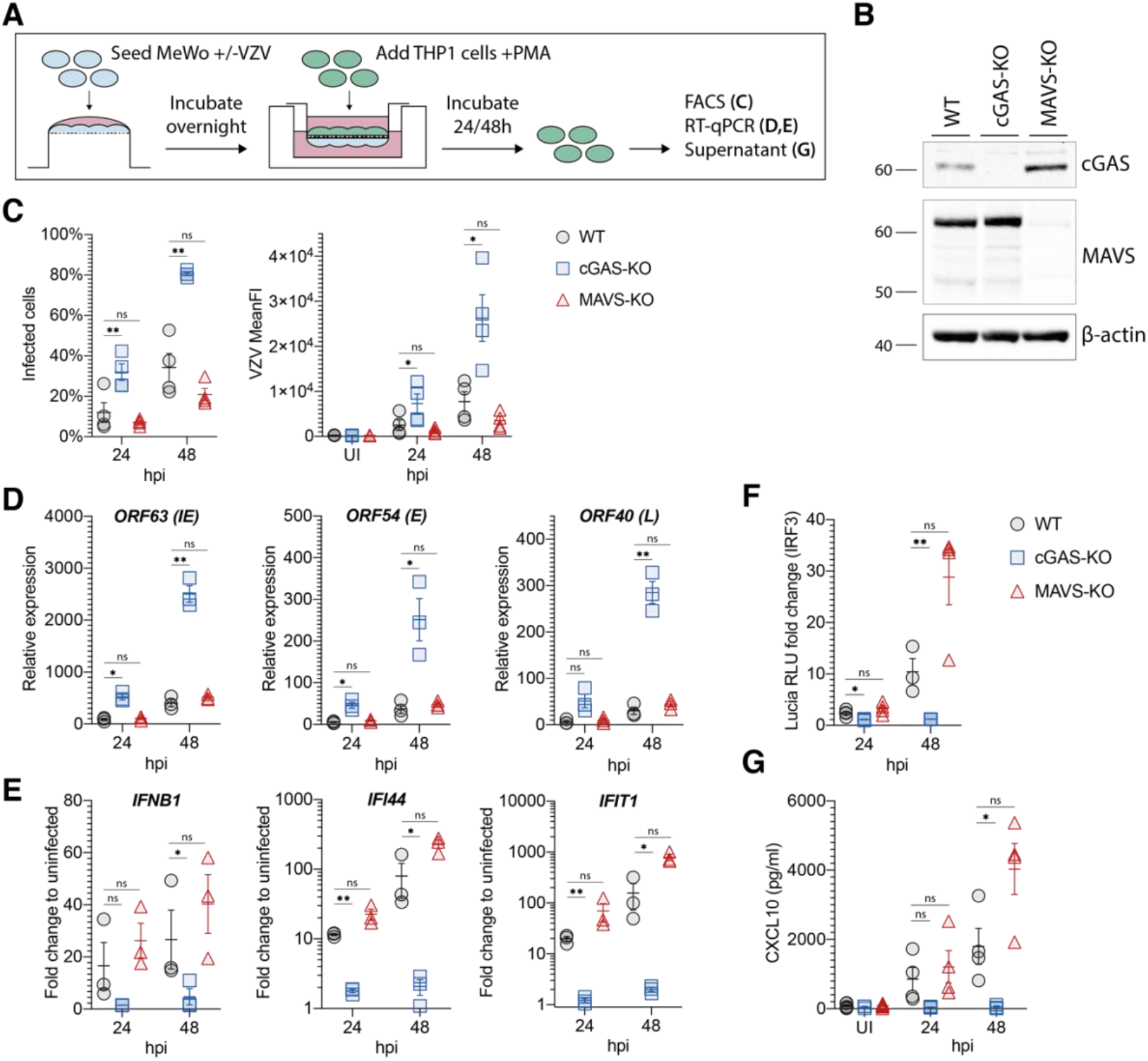
cGAS induces type I IFNs in response to VZV infection. **(A)** Schematic detailing the experimental procedure for infection of PMA-differentiated THP1 cells with VZV by transwell assay. See text for details. **(B)** Immunoblot of WT, cGAS KO, and MAVS KO THP1 cells used in (C-G). **(C)** THP1 cells of the indicated genotypes were mock infected or infected with VZV for 24 and 48 hours using the transwell assay described in (A). Infected cells were quantified by surface staining for VZV-gE/gI and flow cytometry analysis. See Figure S4 for gating. **(D)** RT-qPCR analysis of VZV ORF63 (immediate early (IE) gene), ORF54 (early (E) gene), and ORF40 (late (L) gene) transcripts in cells infected as in (C). **(E)** RT-qPCR analysis of *IFNB1*, *IFI44*, and *IFIT1* expression in cells infected as in (C). **(F)** Activity of Lucia luciferase (secreted under an IRF3-dependent promotor) was determined in supernatants of cells infected as in (C) by QUANTI-Luc assay. **(G)** Concentrations of CXCL10 in supernatants of cells infected as in (C) were determined by ELISA. Panels (C), (F), and (G) show pooled data from four independent biological repeats (n=4 ± SEM). Panels (D) and (E) show pooled data from three independent biological repeats (n=3 ± SEM). Statistical analysis in panels (C), (D), (F), and (G) was paired t-tests and in panel (E) paired ratio t-tests. **=p<0.01, *=p<0.05, UI: uninfected, hpi: hours post infection, WT: wild type, KO: knockout. Panel (B) is representative of two experiments. See also Figures S1, S2, S3 and S4.

WT, cGAS KO, and MAVS KO THP1 (Figure 1B) cells were infected for 24 and 48 hours using this transwell assay. Cell-surface staining for the VZV-gE/gI complex and flow cytometry analysis (Mo et al., 2003) revealed significantly higher levels of infection in cGAS KO cells compared to WT and MAVS KO cells at both time points (Figures 1C and S4). We further found increased expression of immediate early (IE), early (E), and late (L) viral gene products in cells lacking cGAS (Figure 1D). In line with our previous results, THP1 cells failed to upregulate *IFNB1* and ISG expression after VZV infection in the absence of cGAS (Figure 1E). In WT and MAVS KO cells, VZV infection robustly induced secretion of the IRF3-controlled luciferase reporter and CXCL10 (Figure 1F,G). This response was undetectable in cGAS KO cells.

These results confirm our previous observations from the co-culture system and establish that the type I IFN response to VZV infection in THP1 cells was mediated by the DNA sensor cGAS. Significantly more cells became infected with VZV in the absence of cGAS, indicating that recognition by cGAS was required for restriction of VZV infection.

### A VZV ORF Expression Library

Type I IFNs inhibit VZV infection (Kim et al., 2017; Ku et al., 2016). VZV, like many other viruses, employs immune evasion strategies that target the type I IFN system. For example, both ORF61 and ORF62 limit IRF3 activation through distinct mechanisms (Sen et al., 2010; Zhu et al., 2011). In light of our finding that cGAS was crucial for type I IFN induction in response to VZV, we hypothesised that the virus expresses a direct antagonist of cGAS and/or STING. Indeed, other large DNA viruses often encode multiple antagonists of the same innate immune pathway (Smith et al., 2018; Stempel et al., 2019). In order to test the role of individual viral gene products in immune evasion we generated an expression library for all canonical VZV ORFs. All coding sequences were PCR-amplified and cloned into a gateway entry vector. Using recombination, these sequences were then shuttled into a lentiviral vector (pLenti6.3/TO/V5). This vector allows expression with a C-terminal V5 epitope tag either after transient transfection or *via* lentiviral transduction. To validate these constructs, we transiently transfected HEK293T cells and analysed expression of VZV proteins by immunoblot using an antibody against the V5 tag (Figure S5). 59 of 72 constructs (82%) were expressed and bands at the expected molecular weights were detected. An additional five constructs were expressed but not at the expected size, and eight were not expressed at detectable levels. This VZV ORF library is a resource for the scientific community and is available to all interested scientists.

### VZV ORF9 is an Antagonist of DNA sensing

To investigate whether VZV ORFs block cGAS/STING activation, we utilised a luciferase-based screening platform in HEK293T cells. In brief, a plasmid expressing Firefly luciferase under *IFNB1* promotor control and pRL-TK, which constitutively expresses Renilla luciferase, were transiently transfected. HEK293T cells do not express cGAS and STING naturally. We therefore reconstituted human cGAS and human STING by transient transfection, which results in activation of the *IFNB1* promotor and firefly luciferase expression. Lastly, individual viral ORFs (or controls) were co-expressed. Firefly luciferase expression was normalised to Renilla luciferase expression, and we calculated for each ORF a luciferase fold change to an empty vector control condition without cGAS and STING expression constructs. The mean and standard deviation of all data points was then used to calculate Z-values, which represent the number of standard deviations an individual data point is diverging from the mean. We used these Z-scores to rank ORFs in their ability to block IFN activation downstream of cGAS/STING (Figure 2A). KSHV ORF52, a previously described cGAS-antagonist (Wu et al., 2015), and the L protein of EMCV, a previously described IRF3-antagonist (Freundt et al., 2018), served as positive controls. As expected, we found these with the lowest ranks (i.e. smallest fold change) in our assay and both potently blocked firefly luciferase induction. In addition, we identified a number of VZV ORFs that showed similar behaviour. The two aforementioned IRF3 antagonists expressed by VZV, ORF61 and ORF62, were among them, which further validated our approach.

**Figure 2.**
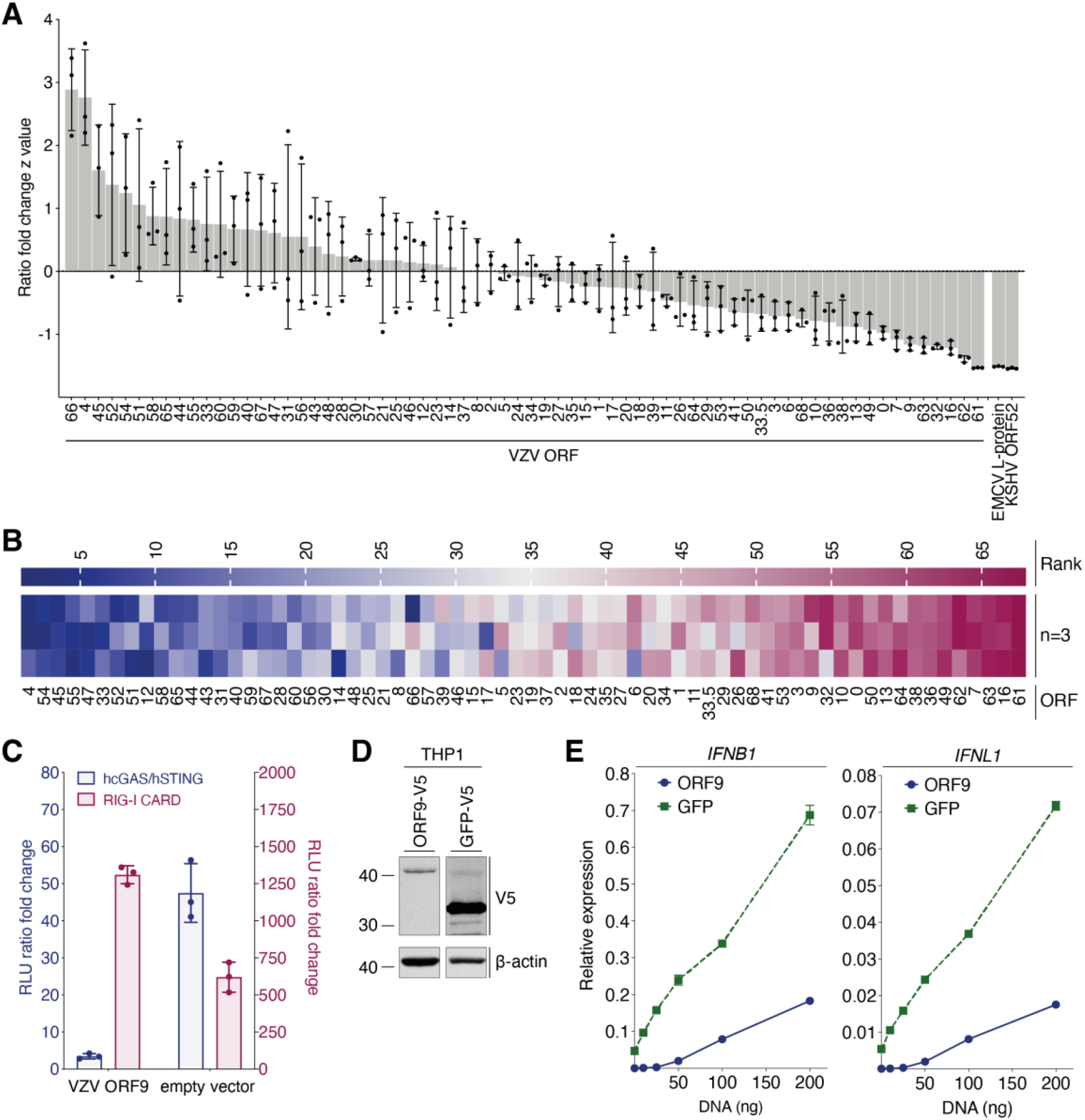
Screening of VZV ORFs identifies ORF9 as an antagonist of cGAS-mediated DNA sensing. **(A)** HEK293T cells were transfected with p125-F-Luc, pRL-TK, expression plasmids for human cGAS and human STING, as well as expression plasmids for individual VZV ORFs. The next day luciferase activity was determined. For each ORF a Z-value was calculated and ORFs were sorted in descending order. **(B)** The experiment shown in (A) was repeated three times and for each ORF in each experiment a rank was determined (highest Z-value = rank 1). Ranks are displayed as a heatmap. The order of the ORFs in the heatmap was determined by their average rank. **(C)** HEK293T cells were transfected as in (A) with expression constructs for VZV ORF9 or empty vector. In parallel, reporter expression was stimulated by co-transfection of the RIG-I-CARD plasmid instead of hcGAS and hSTING. A luciferase fold change was calculated to unstimulated cells. **(D)** Immunoblot analysis of THP1 cells stably transduced with either VZV ORF9-V5 or GFP-V5. **(E)** The same cells as in (D) were differentiated with PMA and transfected with the indicated doses of dsDNA. Expression of *IFNB1* and *IFNL1* mRNAs was assessed by RT-qPCR. Graphs show expression relative to *GAPDH*. Panel (A) is representative of three independent experiments, which are summarised in panel (B). Panels (C-E) are representative of two independent experiments. Data points are technical triplicates with mean and standard deviation (A, C) or the average of technical duplicates with range (E). See also Figure S5.

We performed this screening experiment three times; Figure 2B displays the results as a heatmap. We identified a number of ORFs that reproducibly ranked very low. To identify VZV proteins that specifically block cGAS and/or STING, and not downstream signalling proteins such as IRF3 that are also activated by other PRRs, we compared selected hits from the primary screen in their ability to block reporter activation by overexpression of cGAS/STING or RIG-I-CARD (data not shown). RIG-I-CARD is a constitutively active variant of RIG-I that activates the *IFNB1* promotor *via* MAVS. A direct cGAS/STING antagonist is therefore unable to block this stimulation. We identified the protein encoded by ORF9 of VZV to selectively block reporter activation by cGAS/STING but not RIG-I-CARD when compared to empty vector (Figure 2C).

We then aimed to verify that VZV ORF9 antagonises activation of cGAS by dsDNA in an endogenous setting. THP1 monocytes were stably transduced with ORF9-V5 or GFP-V5 as a negative control (Figure 2D). We stimulated cGAS in these cells by transfection of increasing doses of dsDNA and measured expression levels of *IFNB1* mRNA and *IFNL1* mRNA (encoding a type III IFN) by RT-qPCR (Figure 2E). As expected, THP1 cells expressing GFP showed a dose-dependent increase in expression of both transcripts. In contrast, cells expressing ORF9 did not respond to low doses of DNA. At higher doses, their response was attenuated when compared to GFP-expressing cells. Collectively, these data revealed that the VZV ORF9 protein prevented cGAS/STING activation.

### ORF9 Interacts With cGAS

We hypothesised that VZV ORF9 exerts its antagonistic function by directly interacting with either cGAS or STING. To test this, HEK293T cells were transiently transfected with expression constructs for cGAS-FLAG, STING-HA, and ORF9-V5 and protein interaction was analysed by immunoprecipitation (IP). We used antibodies against the epitope tags and analysed IP fractions by immunoblotting (Figure 3A). All ectopically expressed proteins were precipitated efficiently, and an IgG isotype control antibody did not precipitate any of the proteins tested. Interestingly, ORF9 was detected in the bound fraction after cGAS IP. The reverse IP confirmed this result: we found cGAS in the ORF9 IP. In contrast, ORF9 did not interact with STING in this assay (Figure 3A). To verify that this interaction occurred with endogenous cGAS, we used THP1 cells stably transduced to express FLAG-ORF9, which was precipitated from cell lysates with α-FLAG antibody. Indeed, endogenous cGAS was present in the IP fraction; IP of FLAG-GFP served as a negative control (Figure 3B).

**Figure 3.**
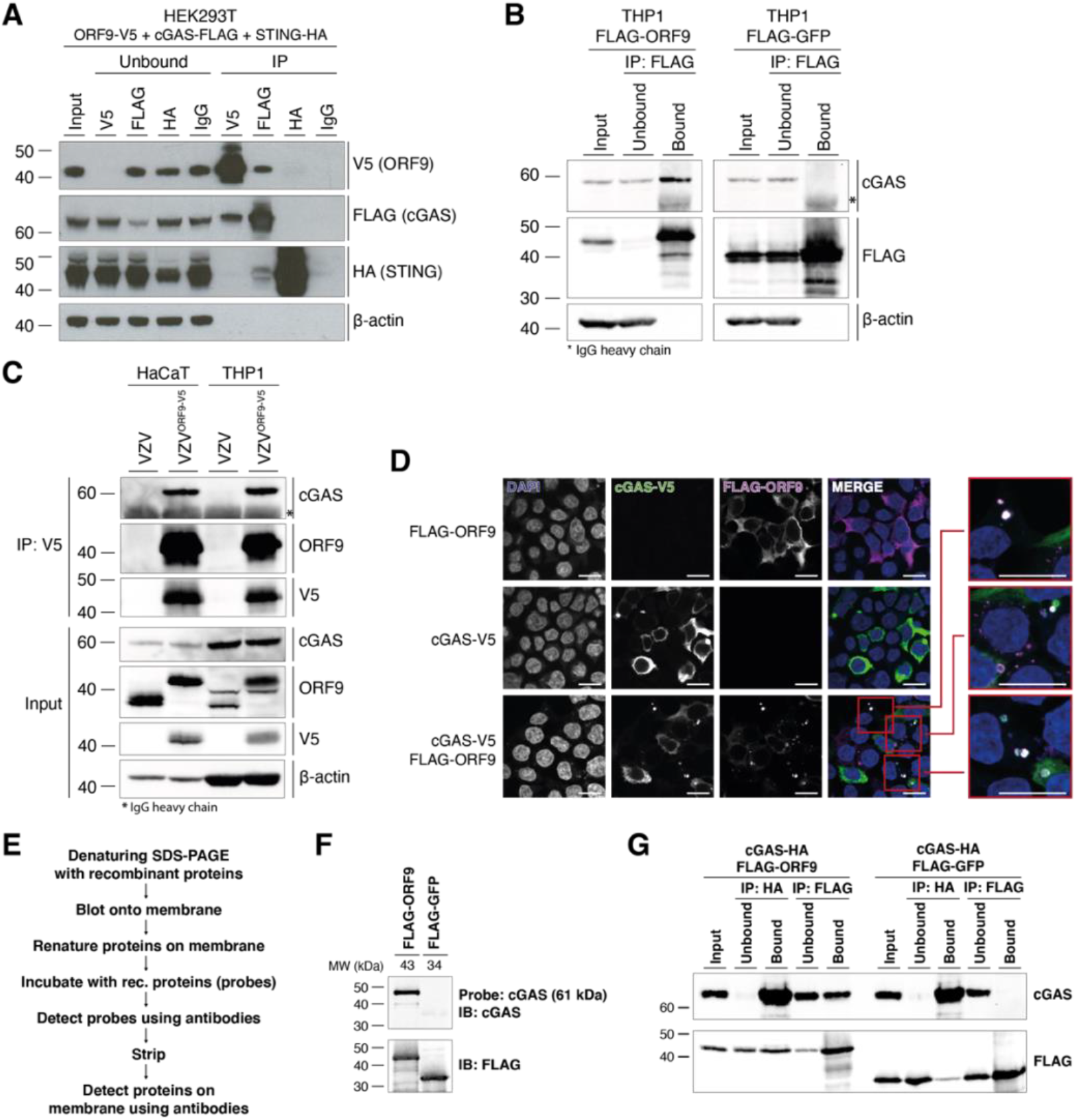
ORF9 interacts with cGAS. **(A)** HEK293T cells were transfected with expression plasmids for ORF9-V5, human cGAS-FLAG, and human STING-HA. The next day, cells were lysed, and overexpressed proteins were immunoprecipitated with α-V5, α-FLAG, α-HA, or IgG isotype control antibody. Input, unbound and IP fractions were subjected to immunoblotting using the indicated antibodies. **(B)** THP1 monocytes stably transduced with either VZV FLAG-ORF9 or FLAG-GFP were PMA-differentiated overnight. The next day, cells were lysed and ectopically expressed proteins were immunoprecipitated using α-FLAG antibody. Input, unbound and IP fractions were subjected to immunoblotting using the indicated antibodies. **(C)** HaCaT cells and PMA-differentiated THP1 cells were infected with WT VZV or VZVORF9-V5 through co-culture with infected MeWo cells for 48 hours. Cells were lysed and ORF9 was immunoprecipitated using α-V5 antibody. Input and IP fractions were subjected to immunoblotting using the indicated antibodies. **(D)** HEK293T cells were seeded onto glass coverslips and were transfected with human cGAS-V5, FLAG-ORF9, or both together. The next day, cells were fixed, permeabilised and stained using α-V5-FITC, rabbit-α-FLAG, and goat-α-rabbit-AF647 antibodies, and DAPI. Mounted coverslips were imaged using confocal microscopy. Scale bars: 15 μm. **(E)** Outline of the far western protocol. **(F)** Far western analysis of cGAS-ORF9 interaction. Recombinant FLAG-ORF9 and FLAG-GFP protein were run on an SDS-PAGE gel and transferred to a membrane. After renaturation of proteins, the membrane was incubated with recombinant human cGAS as a probe, which was detected using α-cGAS antibody. After stripping, the proteins on the membrane were detected using α-FLAG antibody. **(G)** Recombinant human cGAS-HA was mixed with recombinant FLAG-ORF9 or FLAG-GFP. Proteins were immunoprecipitated using α-HA and α-FLAG antibodies. Input, unbound and IP fractions were analysed by immunoblotting. Recombinant proteins used in (F) and (G) are shown in Figure S6A. Panels (A) and (G) are representative of two independent experiments. Panels (B), (D), and (F) are representative of three independent experiments. Panel (C) is representative of two (HaCaT) and three (THP1) independent experiments. See also Figure S6.

Next, we asked whether ORF9 expressed from its endogenous promotor during viral infection had the ability to interact with cGAS. We used a recombinant VZV that expressed C-terminally V5-tagged ORF9 from its endogenous genomic locus. We infected THP1 cells and HaCaT cells (a keratinocyte cell line that expresses cGAS) with WT VZV or VZV^ORF9-V5^ and performed α-V5 IP (Figure 3C). In both THP1 and HaCaT cells infected with VZV^ORF9-V5^, endogenous cGAS co-precipitated with ORF9. These data showed that endogenous ORF9 protein expressed by the virus in infected cells interacted with cGAS.

To investigate whether the two proteins interact in cells, we overexpressed tagged ORF9 and cGAS in HEK293T cells and performed immunofluorescence analysis (Figure 3D). Expression of ORF9 alone resulted in cytoplasmic staining. Similarly, cGAS was detected in the cytoplasm. We further observed cGAS foci co-localising with extranuclear DNA. Extranuclear DNA foci, sometimes in the form of micronuclei, can be observed in some cancer cells and have been shown to bind cGAS (Harding et al., 2017; Hu et al., 2019; Mackenzie et al., 2017). Interestingly, when ORF9 and cGAS were expressed together, both proteins co-localised in DAPI-positive, extranuclear regions. This indicated that ORF9 interacted with cGAS in cells and localised together with cGAS in DNA-positive areas.

In order to biochemically characterise the interaction between ORF9 and cGAS in more detail, we performed experiments in a cell-free system. We expressed cGAS, cGAS-HA, FLAG-ORF9, and FLAG-GFP in *E. coli* and performed single-step purification (Figure S6A). First, we tested whether ORF9 and cGAS interacted directly using the far western protocol (Figure 3E). FLAG-ORF9 or FLAG-GFP protein were separated on a denaturing SDS-PAGE gel and transferred to a membrane. The proteins on the membrane were then re-natured and incubated with recombinant cGAS as a probe. Binding of cGAS to proteins on the membrane was then tested using α-cGAS antibody. Indeed, we found that probing for cGAS resulted in a signal at the size of ORF9 (Figure 3F). Importantly, cGAS did not bind to GFP.

We then tested interaction of the two recombinant proteins using immunoprecipitation. FLAG-ORF9 or FLAG-GFP were incubated with cGAS-HA in the test tube. The proteins were then precipitated using antibodies against the epitope tags. Immunoblot analysis of the IP fractions showed that cGAS was co-immunoprecipitated with ORF9 but not GFP (Figure 3G). The reverse IP of cGAS resulted in binding of both ORF9 and GFP; however, the signal was stronger for ORF9. Taken together, these data indicate that VZV ORF9 and cGAS interacted without the requirement for another cellular or viral protein.

To characterise the interaction of ORF9 and cGAS mechanistically, we constructed ORF9 truncation mutants (Figure 4A). We tested their ability to interact with cGAS by co-IP after overexpression in HEK293T cells (Figure 4B). Consistent with our earlier observation, full-length ORF9 co-immunoprecipitated cGAS. The C-terminal half of ORF9 (construct II) behaved the same, while the N-terminal half (construct I) failed to interact with cGAS. ORF9 constructs III, IV, and V also pulled down cGAS. All ORF9 constructs that interacted with cGAS shared amino acids (AA) 151 to 240. However, IP of this region in isolation (ORF9^151-240^, construct VI) did not co-precipitate cGAS (Figure 4B). Extension of this construct at both ends by about 10 amino acids to generate construct VIII (AA 141 to 249) restored robust interaction with cGAS (Figure 4B).

**Figure 4:**
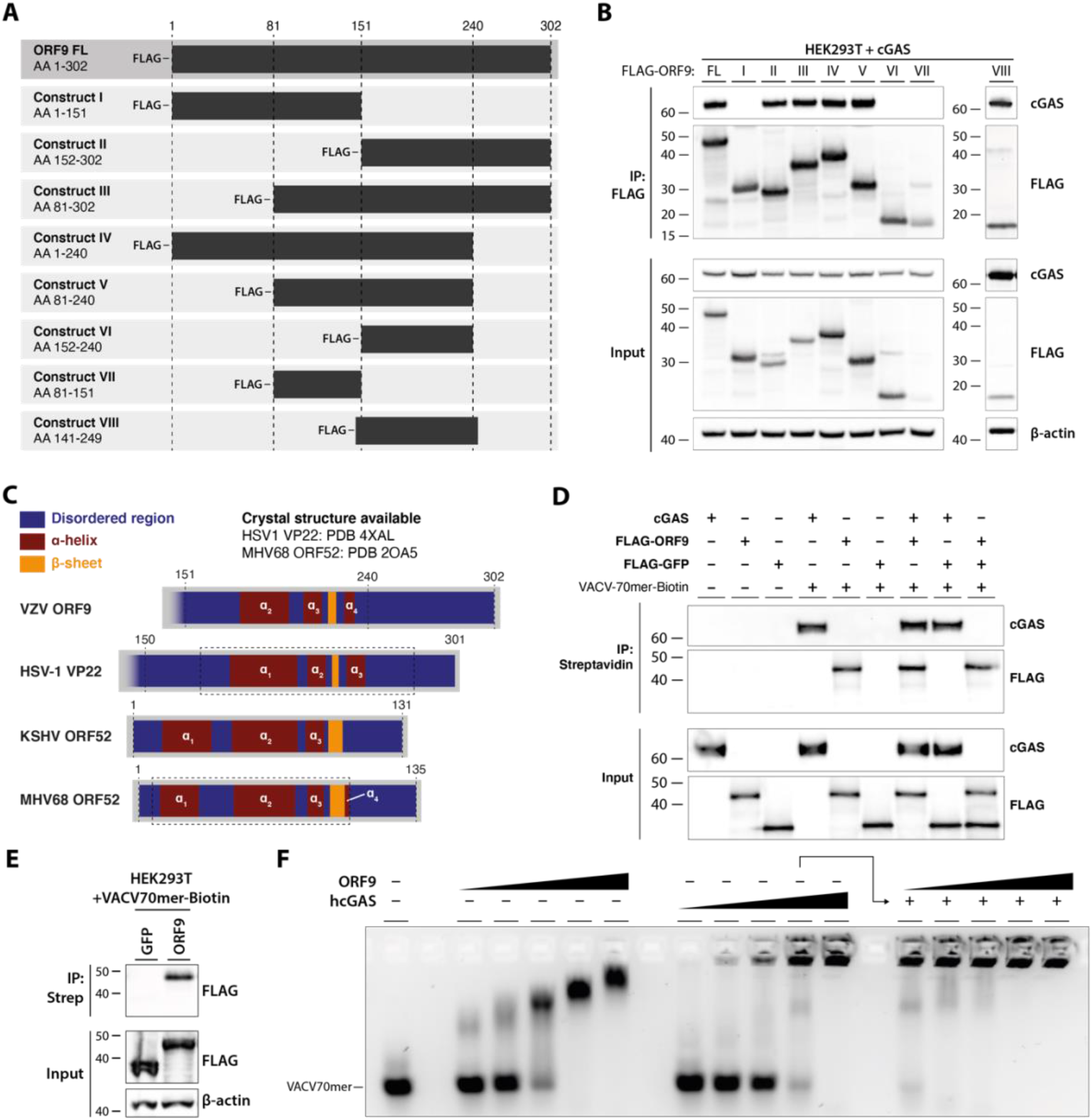
VZV ORF9 binds DNA. **(A)** Schematic detailing ORF9 truncation mutants used in (B). **(B)** HEK293T cells were transfected with expression plasmids for human cGAS and ORF9 truncation mutants. The next day cells were lysed and ORF9 proteins were immunoprecipitated using α-FLAG antibody. IP fractions were subjected to immunoblotting using the indicated antibodies. **(C)** Schematic detailing predicted structural features of VZV ORF9 in relation to predicted and crystal structure-derived features in HSV-1 VP22, KSHV ORF52, and MHV68 ORF52. See text for details. **(D)** Recombinant cGAS, FLAG-ORF9, FLAG-GFP, and biotinylated VACV70mer dsDNA were incubated in the indicated combinations. Lysates were precipitated using streptavidin beads. Fractions were analysed by immunoblotting. **(E)** HEK293T cells were transfected with expression plasmids for FLAG-GFP or FLAG-ORF9. The next day cells were lysed, and the lysate was spiked with biotinylated VACV70mer dsDNA. DNA was precipitated using streptavidin beads and fractions were subjected to immunoblotting using the indicated antibodies. **(F)** VACV70mer dsDNA was incubated with indicated proteins and analysed by agarose gel EMSA. Triangles indicate concentrations of ORF9 (0.7, 1.4, 2.9, 5.7, 8.6 μM), cGAS (0.6, 1.1, 2.2, 4.5, 6.7 μM), and ORF9 (as before) in the presence of 4.5 μM cGAS. Recombinant proteins used in (D) and (F) are described in Figures S6A and S6B, respectively. Results shown in panels (B) and (D-F) are representative of two independent experiments. See also Figure S6.

### ORF9 Binds DNA

To gain insight into possible structural features of ORF9 in this region we used the PSIPRED 4.0 algorithm to predict its secondary structure based on the AA sequence (Buchan and Jones, 2019). This analysis predicted a two helix – sheet – helix motif in the C-terminal region of ORF9 (Figure 4C). We obtained a similar secondary structure prediction for the C-terminal half of VP22, the HSV-1 homologue of VZV ORF9. A crystal structure is available for this region of VP22 (Hew et al., 2015) and confirms the presence of the predicted two helix – sheet – helix motif. Hew et al. further identified a structural similarity of VP22 with the unrelated ORF52 protein of murine herpesvirus 68 (MHV68). MHV68 is closely related to human herpesvirus 8, also known as Kaposi sarcoma associated herpesvirus (KSHV). KSHV ORF52 has been identified as a cGAS antagonist and has DNA-binding properties (Wu et al., 2015). This led us to hypothesise that ORF9 interacts with DNA. To test this, we incubated biotinylated VACV-70mer dsDNA with recombinant ORF9; recombinant cGAS and GFP served as positive and negative controls, respectively. The VACV-70mer is a well-established immunostimulatory dsDNA that binds cGAS (Almine et al., 2017; Lum et al., 2018; Unterholzner et al., 2010). The DNA was then precipitated using streptavidin beads and the presence of bound proteins was analysed by immunoblotting (Figures 4D and S6A). In the absence of DNA, none of the proteins were precipitated. As expected, cGAS bound to DNA. Interestingly, ORF9 was also pulled down by DNA, both alone and in the presence of cGAS. GFP did not bind DNA under any conditions. We further overexpressed either FLAG-ORF9 or FLAG-GFP in HEK293T cells and performed a similar precipitation experiment after spiking the lysates from these cells with VACV70mer-Biotin (Figure 4E). As expected, ORF9, but not GFP, could be pulled down with DNA. To characterise the interaction of ORF9 and DNA in more detail, we performed agarose gel electromobility shift assays (EMSA) (Figures 4F and S6B). Recombinant ORF9 protein impaired the mobility of VACV70mer dsDNA, indicating the formation of ORF9-DNA complexes. These increased in size with higher doses of ORF9, suggesting multivalent protein-protein/protein-DNA interactions. We performed similar experiments with full-length human cGAS protein. As previously described (Zhou et al., 2018), hcGAS and DNA form high molecular weight (HMW) complexes which are unable to migrate out of the gel pocket. Next, we tested incubation of VACV70mer and cGAS with increasing doses of ORF9 protein. Addition of ORF9 to cGAS and DNA increased the size of protein/DNA complexes observed at this concentration of cGAS alone. Taken together, these data show that ORF9 interacted with both DNA and cGAS, without displacing cGAS from DNA.

### ORF9 Phase Separates With DNA

Liquid-liquid phase separation contributes to cGAS activation by dsDNA (Du and Chen, 2018; Xie et al., 2019; Zhou et al., 2021). This is driven by multivalent interactions between cGAS and DNA. In light of our results that ORF9 bound cGAS and DNA, we investigated the effect of ORF9 on cGAS-DNA phase separation. As reported previously, we observed droplet formation by human cGAS and labelled dsDNA, which was sensitive to increasing salt concentration >250 mM (Figures 5A and S6C). Similarly, ORF9 and labelled dsDNA formed liquid droplets in the absence of cGAS (Figures 5A and S6C). ORF9-DNA droplets were smaller than cGAS-DNA droplets, which may indicate a lower propensity of ORF9 to phase separate with DNA compared to cGAS. A C-terminally truncated version of ORF9 (ORF9-N, AA 1–244) had much reduced ability to form liquid droplets (Figures 5A and S6C). Next, we formed cGAS-DNA droplets in the presence of ORF9 and observed that ORF9 had little effect on droplet formation by cGAS and DNA (Figure 5B). Therefore, ORF9 did not antagonise phase separation of cGAS and dsDNA, suggesting another mechanism of cGAS inhibition.

**Figure 5:**
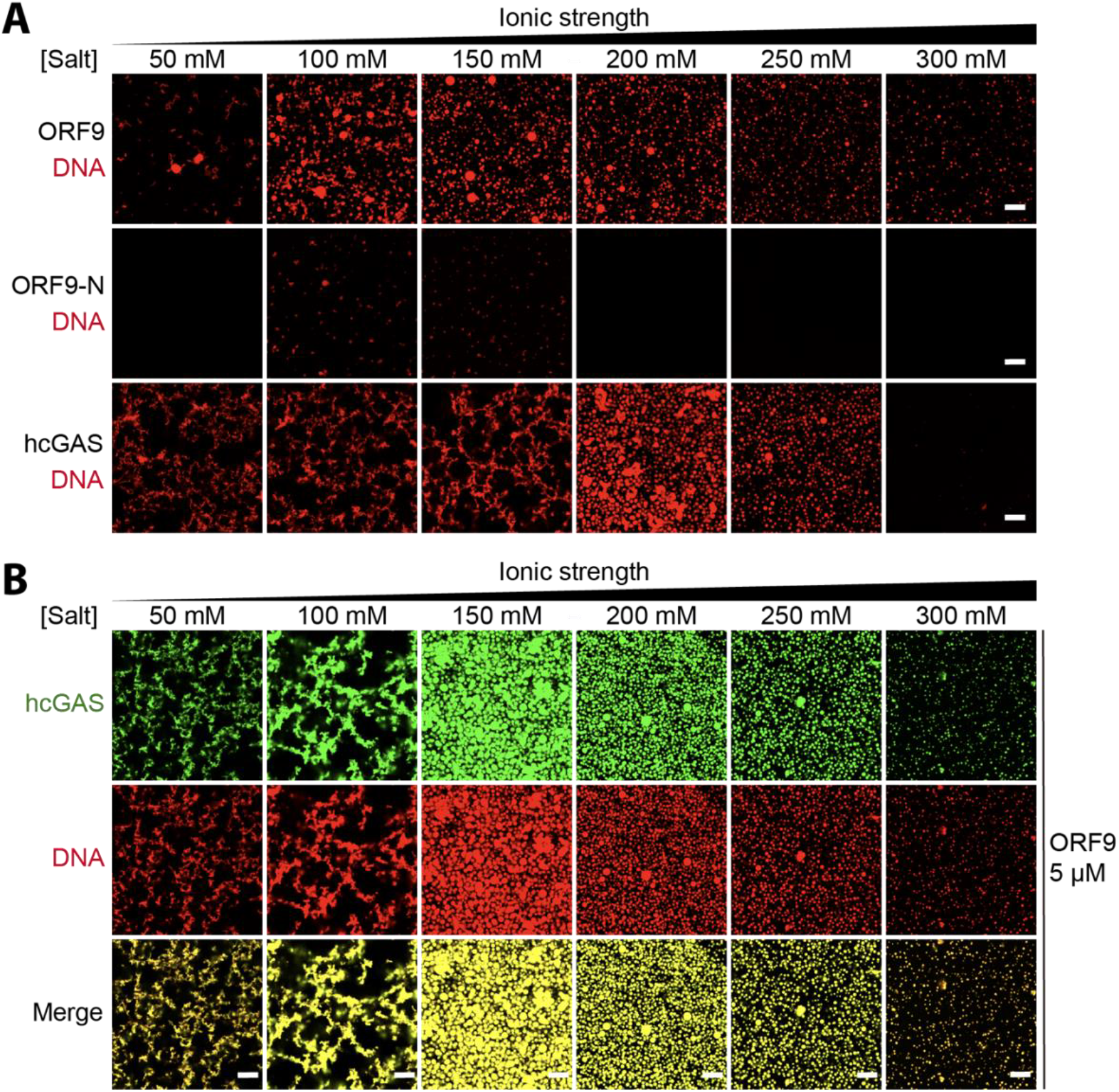
ORF9 phase-separates with DNA. **(A)**10 μM ORF9, ORF9-N, or hcGAS were incubated with 10 μM dsDNA-100 (with 10% of DNA Cy3-labelled) in buffers with increasing salt concentrations (total NaCl and KCl) and imaged by confocal microscopy. **(B)**10 μM hcGAS (with 10% AlexaFluor-488-labelled) was incubated with 10 μM dsDNA-100 (with 10% of DNA Cy3-labelled) and 5 μM ORF9 in buffers with increasing salt concentrations (total NaCl and KCl) and imaged by confocal microscopy. Recombinant proteins used in are described in Figure S6C. Scale bar, 10 μm. Data are representative of 3 experiments. See also Figure S6.

### VZV ORF9 Inhibits cGAMP Synthesis

We next sought to establish the importance of DNA binding for ORF9’s function as a cGAS antagonist. Alignment of the ORF9 with related herpesvirus proteins (Figure 4C) revealed multiple conserved, positively-charged AA residues (Figure 6A). This included ORF9 K178A/R179A and ORF9 R186/R187. The latter aligned with KSHV ORF52 R68/K69 that are required for DNA binding (Wu et al., 2015). We therefore generated three ORF9 mutants: K178A/R179A (construct A), R186A/R187A (construct B), and a double mutant (DM), in which all four residues were mutated to alanine. The mutation sites A and B are located at the beginning and in the middle, respectively, of the first alpha helix in the predicted two helix - sheet - helix of ORF9 (Figure 6B). In the HSV-1 VP22 crystal structure, the corresponding helix forms part of a large, positively charged groove, consistent with a possible role in DNA-binding (Hew et al., 2015). To test the effect of these mutations experimentally, we performed DNA pull-down experiments. As observed before (Figure 4E), WT ORF9 robustly precipitated with DNA (Figure 6C). In contrast, both the A and B mutant were impaired in their ability to bind DNA, with a stronger effect for the B-site. The double mutant did not detectably interact with DNA. Next, we investigated whether the mutant ORF9 proteins interacted with cGAS (Figure 6D). Mutation of either the A or B site alone had no effect on cGAS binding but ORF9-DM showed attenuated cGAS binding.

**Figure 6:**
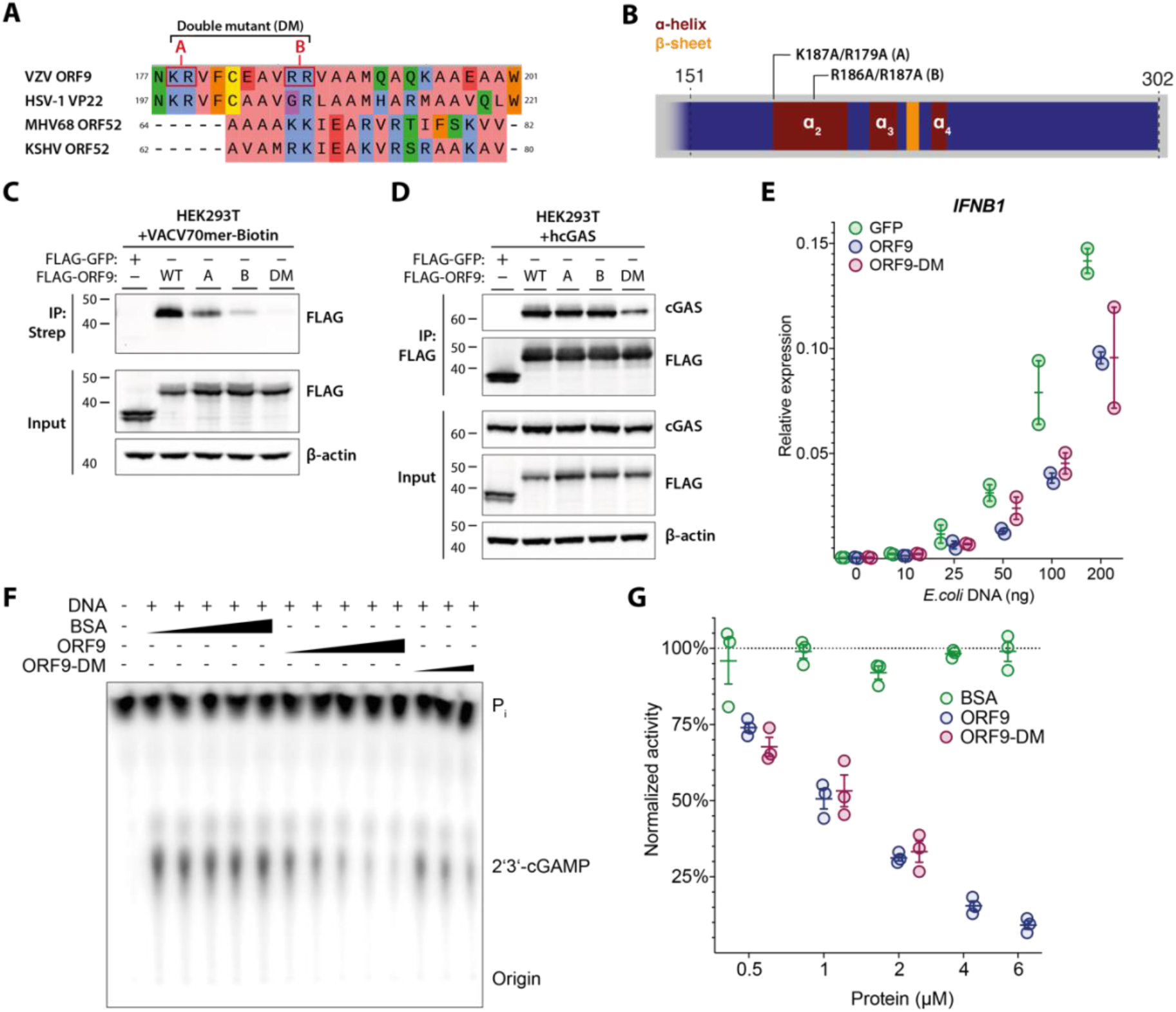
ORF9 inhibits cGAS’ catalytic activity. **(A)** Protein sequence alignment of VZV ORF9, HSV-1 VP22, KSHV ORF52, and MHV68 ORF52. Residues selected for mutagenesis in ORF9 are highlighted in red. Residue colouring represents physico-chemical properties (green: hydrophilic, blue: positive, coral: aliphatic/hydrophilic, orange: aromatic, yellow: cysteine, red: negative, purple: conformationally special). See text for details. **(B)** Schematic detailing position and identity of residues mutated in C-G. **(C)** HEK293T cells were transfected with expression plasmids for FLAG-GFP or FLAG-ORF9 as indicated. The next day cells were lysed, and the lysate was spiked with biotinylated VACV70mer dsDNA. DNA was precipitated using streptavidin beads and proteins were analysed by immunoblotting using the indicated antibodies. **(D)** HEK293T cells were transfected with expression plasmids for human cGAS and FLAG-GFP or FLAG-ORF9. The next day cells were lysed, and proteins were immunoprecipitated using α-FLAG antibody and analysed by immunoblotting using the indicated antibodies. **(E)** THP1 cells stably transduced with GFP, ORF9, or ORF9-DM were transfected with the indicated amounts of *E.coli* dsDNA. The next day, *IFNB1* expression was assessed by RT-qPCR. **(F)***In vitro* cGAS activity assay. Recombinant hcGAS was incubated with ATP, GTP, and radioactive α32P-ATP at 37°C with addition of dsDNA and other proteins as indicated (triangles represent concentrations of 0.5, 1, 2, 4, 6 μM). ORF9-DM was tested only at 0.5, 1, 2 μM. Reactions were treated with calf-intestinal phosphatase and products were analysed by thin-layer chromatography and phosphorimaging. **(G)** Signal intensities from (F) were determined by densitometry analysis and normalized to the average BSA signal. Recombinant proteins used in (F) and (G) are described in Figure S6B. Data shown in panel (C) are representative of two independent experiments. Data shown in panels (D), (E), and (F) are representative of three independent experiments. Panel (G) shows pooled data from three independent experiments (n=2 +/− range). See also Figure S6.

To test the ORF9-DM protein functionally, we generated THP1 cell lines that stably overexpressed GFP, ORF9, or ORF9-DM and stimulated these cells with increasing doses of dsDNA (Figure 6E). Compared to GFP-expressing cells, we found equally reduced *IFNB1* induction in cells expressing either WT or mutant ORF9. These results indicate that the interaction of ORF9 and cGAS was partially dependent on ORF9’s ability to bind DNA. Intriguingly, however, DNA-binding by ORF9 was not required for its ability to inhibit DNA sensing.

Lastly, we used a cell-free cGAMP synthesis assay (Kranzusch et al., 2013) to test whether ORF9 inhibits cGAS enzymatic activity directly. For this, recombinant human cGAS was incubated with radioactively labelled ATP, GTP, DNA, and recombinant ORF9 or bovine serum albumin (BSA) as control. The radioactively-labelled 2′3′-cGAMP produced in these reactions was visualised by thin-layer chromatography and phosphorimaging (Figure 6F and S6B). While addition of BSA did not affect cGAMP production at any dose, ORF9 inhibited cGAS activity in a dose-dependent manner (Figures 6F and 6G). At the doses tested, ORF9-DM was equally potent in inhibiting cGAS compared to WT ORF9.

In sum, these results showed that ORF9 directly inhibited the enzymatic activity of cGAS by a mechanism that did not require DNA binding by ORF9.

## DISCUSSION

VZV is a highly prevalent human virus, yet little is known about its host-pathogen interactions and innate immunity. Here we report that cGAS and its downstream signalling pathway consisting of STING, TBK1 and IRF3 were required for type I IFN induction after VZV infection. An earlier study using RNA interference had implicated STING in innate recognition of VZV (Kim et al., 2017). Our results confirm this observation by complete genetic ablation, and – importantly – identify cGAS as the DNA sensor for VZV. It is possible that innate sensing pathways that detect VZV differ between cell types. Indeed, a study using inhibitory oligonucleotides found that plasmacytoid DCs produce type I IFN in a partially TLR9-dependent manner (Yu et al., 2011). In contrast, we found that the essential TLR9 adaptor protein MyD88 was dispensable for type I IFN induction in VZV-infected THP1 cells. It will be interesting to characterise innate sensing pathways in different cell types relevant to *in vivo* infection, including neuronal cells, T cells, and skin cells. This is important because the viral life cycle and effects of viral replication on host cells can differ between cell types (Zerboni et al., 2014). For example, VZV shows cytopathic effects in fibroblast and some immune cells but does not induce cell death in neurons (Gerada et al., 2020). Future investigation should therefore not only address the role of PRRs in induction of cytokines such as type I IFNs but also in induction of cell death (Maelfait et al., 2020).

Our identification of cGAS as a sensor of VZV infection implicates recognition of an immunostimulatory DNA in infected cells. Studies of herpesvirus entry suggest that the viral DNA remains within the viral capsid during nuclear targeting and may therefore be unavailable for binding to the cytosolic pool of cGAS (Radtke et al., 2006). Single-cell analysis of HSV-1 infection showed that only cells undergoing abortive infection respond by production of type I IFNs (Drayman et al., 2019). It is therefore possible that viral particles with defective capsids are responsible for the type I IFN response observed in bulk populations of cells. Furthermore, cellular restriction mechanisms may make viral DNA from capsids accessible for cGAS binding. Indeed, degradation of herpesviral capsids *via* the ubiquitin – proteasome system has been suggested to release viral DNA for recognition (Horan et al., 2013). Alternatively, cellular DNA may induce type I IFN. Indeed, infections with multiple different viruses result in the accumulation of extranuclear DNA and in mitochondrial damage (Ablasser and Chen, 2019). Host DNA has been shown to stimulate cGAS in the context of HSV-1 infection (West et al., 2015). Whether viral and/or cellular DNA species activate cGAS in VZV-infected cells should be tested in the future by deep sequencing of cGAS-bound DNA. However, this approach has thus far been hampered by the lack of suitable antibodies for specific cGAS IP.

We further describe the identification of ORF9 as a cGAS antagonist. ORF9 interacted with cGAS in a variety of assays, and inhibited cGAMP production and DNA-triggered type I IFN induction. ORF9 is a tegument protein, making it an attractive candidate for antagonising innate immunity. Tegument proteins are contained within viral particles and are delivered into cells at the same time as viral DNA. ORF9 may therefore limit DNA sensing during early infection before viral gene expression. At first glance, our findings that cGAS recognises VZV infection and that ORF9 inhibits cGAS activation might appear contradictory. However, viral immune evasion mechanisms typically limit but not entirely suppress innate immune responses. This notion is supported by the observation that in some cases multiple viral antagonists target the same host defence pathway (Smith et al., 2018; Stempel et al., 2019). In addition, cGAS activation may occur partly in cells infected with defective virions (Drayman et al., 2019) that are likely to contain and/or express reduced amounts of ORF9.

What is the molecular mechanism by which ORF9 inhibits cGAS? Formation of multimeric complexes and higher order structures of cGAS and DNA facilitate cGAS activation (Andreeva et al., 2017; Du and Chen, 2018; Li et al., 2013; Zhang et al., 2014; Zhou et al., 2018). We found that ORF9 binds DNA and phase separated with DNA. We therefore speculated that ORF9 might disrupt cGAS-DNA oligomers. However, further gel shift and phase separation experiments falsified this hypothesis; in contrast, ORF9 might actually facilitate formation of high-molecular weight complexes containing cGAS and DNA. Moreover, an ORF9 mutant with abrogated DNA binding retained the ability to bind cGAS and to inhibit cGAMP production *in vitro* and IFNβ induction *in cellulo*. Although we cannot exclude that this mutant retained weak DNA-binding below the sensitivity of our assays, we prefer a model in which direct protein-protein interactions between ORF9 and cGAS are sufficient for inhibition of cGAMP production. DNA-binding of ORF9 may have unrelated functions and/or may facilitate cGAS interaction and inhibition in specific settings. Future experiments are required to decipher in detail the relationships between ORF9, cGAS, DNA-binding, and cGAMP production. This could include testing DNA length requirements as cGAS oligomer formation is dependent on DNA length (Andreeva et al., 2017; Li et al., 2013; Luecke et al., 2017; Zhou et al., 2018).

ORF9 is essential for viral replication and is a member of the α-herpesvirus UL49 gene family (Che et al., 2008; Tischer et al., 2007). With its closest relative, HSV-1 VP22, it shares 36% AA sequence similarity (Hew et al., 2015). ORF9 has well-established roles in VZV nuclear egress and secondary envelopment (Che et al., 2013; Lebrun et al., 2018; Riva et al., 2013; Riva et al., 2015). Mutational analyses have attributed these functions to AAs or regions either in the N-terminal half or the extreme C-terminus of the protein, whilst we describe a central region (AA151–240) to be required for the interaction of ORF9 with cGAS. A crystal structure is available for the core domain of HSV-1 VP22 that is homologous to this region of ORF9 (Hew et al., 2015). The HSV-1 VP22 core folds into a two helix – sheet – helix motif. Hew et al. further describe structural similarity between HSV1 VP22 and the unrelated MHV68 ORF52 (Hew et al., 2015). MHV68 ORF52 is the homologue of KSHV ORF52. Secondary structure prediction and examination of the published crystal structures revealed that VZV ORF9, HSV1 VP22, KSHV ORF52, and MHV68 ORF52 all potentially share a two helix – sheet structural feature (Figure 4C). Interestingly, both HSV-1 VP22 and KSHV ORF52 have been described previously to inhibit cGAS activation (Huang et al., 2018; Wu et al., 2015). Antagonism of cGAS by KSHV ORF52 requires its DNA binding properties (Wu et al., 2015). In contrast, we found that ORF9’s DNA-binding ability was not required for inhibition of cGAS. This indicates that while structural similarity between the aforementioned proteins could confer cGAS inhibitory properties, the precise molecular mechanisms might differ. In addition, VZV ORF9 and KSHV ORF52 are essential viral gene products while HSV-1 VP22 is not required for viral replication (Huang et al., 2018; Li et al., 2016). Future experiment will be required to shed light on the precise relationship between these viral proteins.

Collectively, our observations lead us to put forward a model in which distantly related herpesviruses have retained within unrelated proteins a structural feature that confers cGAS antagonist properties. Alpha, beta, and gamma-herpesviruses have been estimated to have evolutionarily diverged hundreds of millions of years ago (Brito and Pinney, 2018; Davison, 2002; McGeoch et al., 1995). Anemone species that have diverged from humans more than 500 million years ago harbour cGAS-like enzymes (Kranzusch et al., 2015). This indicates that a common ancestral organism expressed such proteins. It further opens up the possibility that ancient herpes viruses evolved the relevant evasion strategies. We hypothesise that antagonism of cGAS by herpesviruses constitutes an ancient molecular mechanism that evolved long before the advent of antiviral cytokines and IRF3 during evolution.

## AUTHOR CONTRIBUTIONS

JH, RER, and JR conceived the study. JH, WZ, RER, PJK, and JR designed experiments and analysed data. JH, WZ, GF, CC, LC, and TD performed experiments. JH, WZ, and RER developed methodology. JH and JR wrote the manuscript with help from all authors. All authors read and approved the final manuscript.

## ACKNOWLEDGMENTS

The authors thank J Cohen, C Sadzot and S Paludan for VZV stocks, antibodies and cell lines, and P Hublitz and Z Holloway (MRC WIMM Genome Engineering Facility) for their help with CRISPR plasmid generation. The authors further thank C Reis e Sousa, F Zhu, G Ogg, L Unterholzner, A Pichlmair, G Towers, T Fujita, J Haas, and M Reijns for reagents and advice. The authors thank M Brinkmann, J McKeating, and members of the Rehwinkel lab for critical discussion.

## MATERIALS AND METHODS

### Cells

All cells were cultured at 37°C and 5% CO_2_ and checked regularly for mycoplasma contamination. Adherent cells were passaged using Trypsin-EDTA (0.25%) dissociation reagent (Gibco) at appropriate confluence. FCS was obtained from Sigma-Aldrich. HEK293 and HEK293T cells (gifts from Caetano Reis e Sousa, The Francis Crick Institute, UK) were maintained in DMEM (Sigma-Aldrich) containing 4.5 g/L glucose, supplemented with 10% (v/v) FCS and 2mM L-glutamine (Gibco) (DMEM complete). HEK293-ISRE-Firefly luciferase reporter cells (clone 3C11) were described previously (Bridgeman et al., 2015). HaCaT cells were a gift from Leonie Unterholzner (Lancaster University, UK) and were maintained in DMEM complete medium. MeWo cells were a gift from Graham Ogg (University of Oxford, UK) and were maintained in MEM supplemented with 10% (v/v) FCS, 2mM L-glutamine (Gibco), 1x Non-essential amino acids (Gibco), and 1mM sodium pyruvate (Gibco). THP1 Dual cells (WT, MyD88-KO, and IFNAR2-KO) were from Invivogen. STING-KO and TBK1-KO THP1 cells were a gift from Soren Paludan (Aarhus University, Denmark) (Holm et al., 2016). MAVS-KO, cGAS-KO, and IRF3-KO cells were generated as described below. All THP1 cell lines were maintained in RPMI (Sigma Aldrich) supplemented with 10% (v/v) FCS and 2mM L-glutamine (Gibco).

### Viruses

VZV rOka (cosmid-derived recombinant pOka) was a gift from Jeffrey Cohen (NIH, Bethesda, USA) (Cohen and Seidel, 1993). The virus was maintained in monolayers of MeWo cells. In brief, monolayers of infected cells were monitored microscopically for cytopathic effect. Cells that showed high level of infection were detached and infected cells were mixed at appropriate ratios (1:2–1:4) with uninfected cells and re-plated. Aliquots of infected cells were cryopreserved in freezing medium (90% FCS, 10% DMSO), stored in liquid nitrogen, and thawed for experiments. VZV^ORF9-V5^ is a recombinant virus, in which a C-terminal V5 tag was added to the coding sequence of ORF9. The virus was a kind gift from Catherine Sadzot (University of Liege, Belgium) (Riva et al., 2013).

### Plasmids

The p125-Luc plasmid (*hIFNB1* promotor firefly luciferase) was a gift from T. Fujita (Kyoto University, Japan) (Yoneyama et al., 2004). pRL-TK was from Promega. RIG-I-CARD was a gift from Andreas Pichlmair (Technical University Munich, Germany). pCMV-VSV-G was a gift from Bob Weinberg (Addgene number 8454). The lentiviral packaging plasmid p8.91 was a gift from Greg Towers (University College London). pGag-eGFP was from the NIH AIDS reagent programme (number 11468). pX458/Ruby/MAVS was described previously (Hertzog et al., 2018). pCMV-3tag-KSHV-ORF52 was a kind gift from Fanxiu Zhu (Florida State University, USA). pX458/Ruby/IRF3 was created as described for the MAVS-targeting plasmid. A sgRNA targeting exon 3 of the IRF3 locus was designed using the MIT algorithm (crispr.mit.edu) and cloned into the px458 vector. The pLenti6.3/EMCV-L-V5 plasmid was described previously (Hertzog et al., 2018).

### Cloning

Mammalian expression plasmids for hcGAS and hSTING were created by PCR-amplification using THP1 cDNA and ligation into the gateway entry vector pCR8. Coding sequences were shuttled into expression vectors (pLenti6.3/C-V5, pcDNA3.2/C-FLAG, pcDNA3.2/C-HA) via Gateway recombination. Mammalian expression plasmids for eGFP (pLenti6.3/puro/N-FLAG, pDEST31/N-FLAG) were created in a similar way by PCR amplification from pGag-eGFP introducing a stop codon. VZV ORF9 expression plasmids (pLenti6.3/puro/N-FLAG, pDEST31/N-FLAG) were created by amplification of ORF9 with a stop codon and gateway recombination. Expression plasmids for ORF9 truncation mutants (pLenti6.3/puro/N-FLAG) were created by PCR amplification of the respective coding sequence with a start and stop codon. ORF9 mutant plasmids were generated using site directed mutagenesis (QuikChange II Site-Directed Mutagenesis Kit, Agilent). GST-fusion bacterial expression vectors for FLAG-ORF9, FLAG-GFP, and hcGAS-HA were created by PCR-amplification of the coding sequences from existing plasmids and restriction enzyme cloning into pGEX6P1. pGEX6P1 and pGEX6P1-hcGAS were a kind gift from Martin Reijns (University of Edinburgh, UK). Further bacterial expression vectors for FLAG-ORF9 and FLAG-ORF9-DM were created by restriction enzyme cloning of coding sequences into the pET28a bacterial expression vector (kind gift from Simon Davis, University of Oxford). Bacterial expression vectors for ORF9 and ORF9-N were created by restriction enzyme cloning of coding sequences into a custom 6xHis-SUMO bacterial expression vector (Zhou et al., 2018). All primer sequences are listed in Table S1.

### Immunostimulatory dsDNA

*E. coli* dsDNA was from Invivogen. A 70bp immunostimulatory dsDNA fragment from VACV was described previously (Unterholzner et al., 2010). Two complementary oligos were synthesised (Sigma Aldrich) and combined at equal molar ratio. The solution was heated to 95°C and allowed to anneal by cooling to RT.

Forward sequence: 5′-ccatcagaaagaggtttaatatttttgtgagaccatcgaagagagaaagagataaaacttttttacgact-3′-TEG-Biotin

Reverse sequence: 5′-agtcgtaaaaaagttttatctctttctctcttcgatggtctcacaaaaa-tattaaacctctttctgatgg-3′

The 100bp dsDNA used for the phase separation assay was described previously (Zhou et al., 2021). DNA oligos were synthesised (Integrated DNA Technologies) and double-stranded DNA was prepared by annealing two complementary oligos.

Forward sequence: 5′-ACATCTAGTACATGTCTAGTCAGTATCTAGTGATTATCTAGACATACATCTAGTACATGTCTAGTCAGTATCTAGTGATTATCTAGACATGGACTCATCC-3′

Reverse sequence: 5′-GGATGAGTCCATGTCTAGATAATCACTAGATACTGACTAGACATGTACTAGATGTATGTCTAGATAATCACTAGATACTGACTAGACATGTACTAGATGT-3′

### Antibodies

<u>For immunoblot</u>: beta-actin-HRP (AC-15, Sigma Aldrich), FLAG-HRP (clone M2, Sigma Aldrich), V5-HRP (R961-25, Invitrogen), RIG-I (Alme1, Calteg Medsystems), VZV gE/GI (MAB8612, GE Healthcare), VZV ORF62 (C05107MA, Meridian Life Science), pSTAT1 (Y701) (58D6, CellSignaling Technology), STAT1 (42H3, CellSignaling Technology), pSTAT2 (D3P2P, CellSignaling Technology), STAT2 (D9J7L, CellSignaling Technology), hcGAS (D1D3G, CellSignaling Technology), hSTING (D2P2F, CellSignaling Technology), MyD88 (D80F5, CellSignaling Technology), TBK1 (D1B4, CellSignaling Technology), MAVS (ALX-210-929-C100, ENZO Life Science), IRF3 (D6I4C, CellSignaling Technology), VZV ORF9 (polyclonal rabbit serum, kind gift from Catherine Sadzot (University of Liege, Belgium) (Riva et al., 2013), donkey-anti-mouse-HRP (NA931, GE Healthcare), donkey-anti-rabbit-HRP (NA931, GE Healthcare). <u>For IP</u>: FLAG (clone M2, Sigma Aldrich, V5 (680602, Biolegend), HA (2-2.2.14, Invitrogen). <u>For IF</u>: V5-FITC (R963-25, Invitrogen), FLAG (D6W5B, CellSignaling Technology), goat-anti-rabbit-AF647 (A21246, Invitrogen). <u>For FACS</u>: VZV-gE/gI (see above) was conjugated to FITC using FITC Conjugation Kit (Abcam).

### VZV ORF Library

Primers for PCR amplification of individual VZV ORF sequences were designed based on an annotated VZV genome sequence (GenBank accession: AB097933.1). Forward primers included a Kozak sequence (GCCGCC), added before the start codon of an ORF. Reverse primers excluded the Stop codon to allow for addition of a C-terminal tag through the vector (see below). Primer sequences are listed in Table S2. The PCR template was generated by extracting RNA from VZV infected MeWo cells using QIAshredder (Qiagen) and RNeasy Mini Kit (Qiagen). The RNA was reverse transcribed into cDNA using SuperScript II Reverse Transcriptase (Invitrogen). For some ORFs, DNA from infected MeWo cells extracted with DNeasy Blood and Tissue Kit (Qiagen) served as the PCR template. PCR products were generated using Phusion High-Fidelity DNA Polymerase (New England Biolabs) or Herculase II Fusion DNA Polymerase (Agilent Technologies). PCR reactions were analysed by agarose gel electrophoresis and PCR products of the predicted size were extracted from the gel using QIAquick Gel Extraction Kit (Qiagen). Fragments were ligated into the pCR8/TOPO Gateway entry vector (Invitrogen). Plasmid DNA from single clones of transformed *E.coli* (New England Biolabs) was analysed for correct orientation of the insert by Sanger sequencing. Inserts were then shuttled into pLenti6.3/TO/V5-DEST (Invitrogen) using LR Clonase II Enzyme Mix (Invitrogen) for Gateway recombination. All clones with the pLenti6.3/TO/V5 backbone were propagated in recombination-deficient Stbl3 bacteria (Invitrogen).

Cloning of the entire coding sequence of ORF22 (8256 bp) was unsuccessful using various amplification and cloning technologies. Therefore, two Gateway entry vectors encoding individual segments within the ORF22 coding sequence (ORF22A: nt 2219-4029, ORF22B: nt 4012–6114, described previously (Uetz et al., 2006)) were used (kind gift from Jurgen Haas, University of Edinburgh, UK). Expression vectors with the pLenti6.3/TO/V5 backbone were generated as described above.

To validate expression, HEK293T were seeded at 5×10^5^ cells per well in 12-well plates. The next day, cells were transfected with 600ng VZV ORF or control expression plasmids using 3μl Lipofectamine 3000 (Invitrogen) per well. 24 hours later, cells were lysed in RIPA buffer (10mM TRIS-HCl pH8, 140mM NaCl, 1% Triton-X 100, 0.1% SDS, 0.1% sodium deoxycholate, 1mM EDTA, 0.5mM EGTA). Lysates were clarified by centrifugation and 30μg protein was subjected to immunoblotting.

### Luciferase reporter assays

HEK293T cells were seeded at 3.5×10^4^ cells per well in 96-well plates. On the following day, cells were transfected with the following plasmids using Lipofectamine 2000 (Invitrogen): 20ng p125-F-Luc, 5ng pRL-TK, 1ng hcGAS, 25ng hSTING, and 50ng of a VZV ORF. Alternatively, cells were transfected with 5ng RIG-I-CARD plasmid instead of cGAS and STING plasmids. The next day, expression of firefly and renilla luciferases was assessed using Dual Luciferase assay system (Promega).

Activity of the secreted Lucia luciferase under IRF3 promoter control (THP1 Dual cells) was assessed using QuantiLuc substrate (Invivogen) according to manufacturer’s instructions.

### Lentivirus Production and Transduction

1.2×10^7^ HEK293T cells were seeded in 15cm cell culture dishes. The next day, cells were transfected with 9μg lentiviral expression plasmids harbouring the gene of interest, 9μg p8.91 plasmid, and 3μg pVSV-G using Lipofectamine 2000. 16 hours later, medium was replaced with fresh growth medium. 24 hours later, lentivirus containing supernatant was harvested, clarified by centrifugation, and stored at 4°C. Cells were overlaid with fresh growth medium. 8 hours later, supernatants were harvested again and pooled with previous supernatants. After 16 hours, lentivirus containing supernatants were harvested for a third time. Pooled supernatants from all three harvests were filtered through a 0.45μm filter, aliquoted into cryovials and stored at −80°C. For transduction of cells, polybrene (Sigma Aldrich) was added to lentiviral supernatants to a final concentration of 8μg/ml. THP1 cells were pelleted and resuspended in lentiviral supernatant containing polybrene. The next day, cells were pelleted and resuspended in fresh growth medium. After overnight incubation, cells were once again pelleted and resuspended in growth medium containing 10μg/ml blasticidin or 1μg/ml puromycin (both Gibco) depending on which vector was used. Surviving cells were used for experiments.

### THP1 Knockout Cell Generation

For generation of knockout cells using CRISPR/Cas9 technology the pX458-mRuby plasmid encoding the Cas9 protein, the sgRNA, and mRuby was used (Hertzog et al., 2018). 1×10^7^ THP1 cells were transiently transfected with 50μg of respective pX458 plasmids using Lipofectamine LTX (Invitrogen) and incubated overnight. The next day cells were stained with 10μg/ml DAPI in PBS for 10min. After resuspension of cells in growth medium, live, mRuby-positive cells were FACS-sorted into eppendorf tubes containing growth medium with 20% FCS. The suspension of sorted cells was diluted with special growth medium (50% conditioned THP1 medium, 40% fresh RPMI with 10% FCS, 10% additional FCS) to one or three cells per 200μl. 200μl cell suspension were then dispensed into each well of a 96-well plate (one or three cells per well). The cells were incubated for several weeks until clones grew out.

The absence of the targeted protein in each clone was assessed by immunoblot analysis. For functional validation of MAVS and IRF3 knockout cells, 1.5×10^5^ cells were seeded per well into 96-well plates in growth medium containing 10ng/ml PMA (Sigma-Aldrich). The next day cells were transfected with 5ng IVT-RNA (Rehwinkel et al., 2010) or 15ng *E.coli* DNA (Invivogen) using Lipofectamine 2000 (Invitrogen). 24 hours later, IFN in supernatants was measured using the type I IFN bioassay. For functional validation of cGAS knockout cells, cells were stimulated in the same way and activity of Lucia luciferase (under IRF3 promoter control) was assessed in supernatants. Knockout clones were further analysed for insertions and deletions in their genomic loci. *MAVS* and *IRF3* target regions were PCR-amplified using Herculase II Fusion DNA Polymerase. PCR amplicons were gel extracted (QIAquick Gel Extraction Kit) and sequenced by Sanger sequencing. Sequencing traces were analysed using the TIDE software (https://tide.nki.nl) (Brinkman et al., 2014).

### VZV Flow Cytometry Infection Assay

Cells were washed in FACS tubes with PBS and incubated with LIVE/DEAD Fixable Violet Dead Cell Stain (Invitrogen) diluted 1:1000 in PBS for 30min at 4°C. Cells were washed in PBS and resuspended in 100μl FACS buffer (1% FCS, 2mM EDTA in PBS) containing 1:500 FITC-coupled antibody against VZV gE/gI complex. Cells were incubated 30min at 4°C. After washing with PBS cells were resuspended in PBS and an equal volume of 8% methanol-free formaldehyde in PBS was added. Cells were fixed for 15min at room-temperature. Cells were washed with PBS, resuspended in FACS buffer, and analysed on a Attune NxT Acoustic Focusing Cytometer (Thermo Fisher).

### THP1 VZV co-culture Infections

6.125×10^6^ THP1 cells were seeded per well in 6-well plates in growth medium containing 10 ng/ml PMA (Sigma-Aldrich). As a control, 1.625×10^6^ MeWo cells were seeded in MEM. Aliquots of uninfected and VZV-infected MeWo cells were thawed, washed, and resuspended in MEM. Cells were counted and their concentration was adjusted to 6.25×10^5^ live cells per ml (for THP1 cells) or 1.25×10^5^ live cells per ml (for MeWo cells) with MEM. Medium was removed from labelled cells and cells were overlaid with 2ml MeWo −/+VZV cell suspension. Cells were incubated at 37°C for 1 hour. The cell suspension was removed, adherent cells were washed with PBS, overlaid with their respective growth medium and incubated for 48 hours. Supernatants were removed from cells, clarified by centrifugation and stored at −80°C. CXCL10 levels were determined using Human CXCL10/IP-10 Quantikine ELISA Kit (R&D Systems) according to manufacturer’s instructions. Cells were washed with PBS and incubated with Trypsin-EDTA until they started to detach. Growth medium was added, cells were resuspended by gentle pipetting and transferred to tubes. Half of the cell suspension were transferred to eppendorf tubes on ice for extraction of RNA and generation of protein lysates, respectively. The cells in those eppendorf tubes were pelleted, washed with PBS, and then lysed either in RLT buffer and processed for RT-qPCR or lysed in RIPA buffer and processed for immunoblot analysis.

### THP1 transwell VZV infections

PET membrane 1 μm transwell inserts (Sarstedt 83.3930.101) were placed with the bottom membrane facing upwards into 15 cm dishes. Aliquots of uninfected and VZV-infected MeWo cells were thawed, washed, and resuspended in MEM. Cells were counted and their concentration was adjusted to 3×10^6^ live cells per ml with MEM. 500 μl cell suspension (1.5×10^6^ cells) was pipetted onto the transwell membrane and cells were let adhere in the incubator overnight. 6-well plates were filled with RPMI media and prepared transwells were placed cell-side downwards into the plates, so that the MeWo cells were submerged. 2×10^6^ THP1 cells were seeded onto the upward facing side of the transwell membrane in RPMI medium containing 10 ng/ml PMA. Cells were harvested by trypsinisation after 24h and 48h and used for downstream analysis. A publication describing this method in more detail is in preparation.

### Pulldowns

1.4×10^7^ HEK293T cells were seeded in 15cm cell culture dishes. The next day, cells were transfected with 12.7μg of ORF9-V5, cGAS-FLAG, and STING-HA expression plasmids using Lipofectamine 2000. 16 hours later, cells were lysed in IP buffer (20mM TRIS-HCl pH7.4, 100mM NaCl, 1mM EDTA, 0.5% NP-40, Protease Inhibitor Cocktail (CellSignaling Technology)). After clarification, an aliquot was removed as input sample and the lysate was split into four equal volumes. Each aliquot was incubated with 50μl Dynabeads Protein G that were coated with 5μg α-V5, α-FLAG, α-HA, or control IgG antibody for 1 hour under rotation at 4°C. The supernatant was removed, an aliquot was stored as unbound fraction from each sample, and beads were washed three times in lysis buffer. Input, unbound, and bound fractions were subjected to immunoblotting. For pulldowns from stably transduced THP1 cells, cells were seeded at 2×10^7^ per dish in 15cm cell-culture dishes in growth medium containing 10ng/ml PMA. Lysates were generated the next day and processed as described above. For pulldowns of recombinant proteins, 3μl recombinant cGAS-HA protein was mixed with 1.5μl FLAG-ORF9 or FLAG-GFP in IP buffer. IP was performed as described above. For pulldowns from VZV-infected cells, 2.2×10^7^ THP1 cells and 4×10^6^ HaCaT cells were seeded in 10cm dishes (in the presence of 10ng/ml PMA for THP1 cells). The next day cells were overlaid with 1.25×10^7^ MeWo cells infected with VZV or VZV^ORF9-V5^ for 1 hour and afterwards washed with PBS. Lysates were generated 48 hours later as described above.

### Far western

The experimental procedure for far western was based on a protocol previously described (Wu et al., 2007). 1.5μl FLAG-ORF9 protein or 1.5μl FLAG-GFP protein was subjected to SDS-PAGE and blotting as described for regular immunoblotting (see below). After transfer, the membrane was incubated for 30min at RT temperature in AC buffer (6M Guanidine HCl, 100mM NaCl, 20mM TRIS-HCl pH 7.6, 0.5mM EDTA, 10% glycerol, 0.1% Tween-20, 2% skim milk powder, 1mM DTT). The membrane was then incubated in AC buffers with decreasing concentrations of Guanidine HCl (3M, 1M, 0.1M) for 30min each. For the last incubation the membrane was transferred to 4°C and thereafter incubated in AC buffer free of Guanidine HCl overnight. The membrane was blocked for 1 hour in 5% milk powder in PBST (PBS with 0.05% Tween-20) and then incubated with 10μl recombinant cGAS per 5ml buffer as probe in 3% milk powder in PBST for 1 hour. Membranes were washed for 10min with PBST thrice and then incubated with α-cGAS antibody for 1 hour. After washing, membranes were incubated with appropriate secondary antibodies, washed again, and imaged. Hereafter, membranes were stripped and re-probed with α-FLAG antibody as described for conventional immunoblot.

### Immunofluorescence

1.75×10^5^ HEK293T cells were seeded onto-glass coverslips. The next day, cells were transfected with 250ng of cGAS-V5 or FLAG-ORF9 expression plasmid using 1.5μl Lipofectamine 3000 per well. Additional coverslips were co-transfected with both plasmids. 24 hours later cells were washed with PBS and fixed with 4% formaldehyde in cytoskeleton stabilisation buffer (CSB; 10mM KCl, 274mM NaCl, 8mM NaHCO_3_, 0.8mM KH_2_PO_4_, 2.2mM Na_2_HPO_4_, 4mM MgCl_2_, 10mM PIPES, 4mM EGTA, 11mM glucose) for 15min. Cells were washed and permeabilised with 0.1% Triton X-100 in CSB for 20min and incubated with 100mM glycine in CSB for 10min afterwards. Cells were washed with PBS four times and blocked using 1% BSA and 5% normal goat serum (Invitrogen) in PBS for 1 hour. Coverslips were incubated with primary antibodies in blocking solution for 3 hours at room temperature. Cells were washed three times with PBS and incubated with secondary antibodies in blocking solution for 1 hour. Coverslips were washed with PBS three times and mounted onto glass slides using ProLong Diamond Antifade Mountant with DAPI (Invitrogen). Images were acquired using a Zeiss LSM780 confocal microscopy system (Zeiss).

### Recombinant Protein Expression

#### Single-step purification of GST-fusion proteins (see Figure S6A)

BL21-pLysS-Rosetta *E.coli* were transformed with bacterial expression vectors. A single colony or a glycerol stock from a single colony were used to inoculate a starter culture of 25ml LB-medium containing 34μg/ml Chloramphenicol and 100μg/ml Carbenicillin. A large 400ml LB culture was inoculated using the starter culture. OD600 was measured in intervals and, for VZV ORF9 and GFP, recombinant protein expression was induced by adding 0.1mM IPTG once OD reached 0.6. Bacteria were grown 3 hours at 37°C. For expression of recombinant cGAS and cGAS-HA, bacteria were grown to OD 0.8 and chilled to 18°C. Protein expression was induced by adding 0.4mM IPTG and bacteria were incubated for 16 hours at 18°C. For all proteins, bacteria were pelleted at 6,000g for 7min and the pellet was resuspended in 12ml lysis buffer (20mM TRIS pH7.4, 500mM NaCl, 0.5mM EDTA, 0.5mM EGTA, 0.5% NP40, 1:100 Protease Inhibitor Cocktail (CellSignaling Technology)). The suspension was sonicated on ice three times at 20% amplitude with 15 sec ON and 30 sec OFF (Branson Sonifier). Insoluble material was removed by centrifugation for 15min at 24,000g. In the meantime, 150μl Glutathione Sepharose beads (GE Healthcare) were prepared according to manufacturer’s instructions. The beads were added to the clarified lysate and incubated under rotation for 4 hours at 4°C. The beads were pelleted by centrifugation at 500g and washed five times with lysis buffer. The beads were washed once in PreScission Cleavage buffer (50mM TRIS pH 7.5, 150mM, 1mM freshly added DTT) and resuspended in 500μl PreScission Cleavage buffer. After addition of 12μl PreScission Protease (GE Healthcare), the suspension was incubated under rotation for 3 hours at 4°C. The supernatant containing the recombinant protein was separated from beads and stored at −80°C. Aliquots from the various purification steps were analysed by SDS-PAGE and subsequent staining of the gel with EZBlue Gel Staining Reagent (Sigma Aldrich). Based on band intensities the protein concentrations were estimated to be ca. 1μg/μl for FLAG-ORF9 and FLAG-GFP, and ca. 0.5μg/μl for cGAS and cGAS-HA.

#### ORF9 purifications for EMSA and cGAS activity assay (Figure S6B)

BL21-pLysS-Rosetta *E.coli* (Novagene) were transformed with bacterial expression vectors. Three fresh, single colonies were used to inoculate a starter culture of 150ml LB-medium containing 34μg/ml chloramphenicol and 50μg/ml kanamycin and grown overnight at 16°C. Two large 1 L LB cultures were then inoculated using the starter culture. Bacteria were grown to OD 0.8 and chilled to 16°C. Protein expression was induced by adding 0.5mM IPTG and bacteria were incubated for 16 hours at 16°C. Bacteria were pelleted at 6,000 g for 15 min and the pellet was resuspended in 120 ml lysis buffer (20 mM HEPES-KOH pH 8, 400 mM NaCl, 0.5% NP40, 10% glycerol, 30 mM imidazole, 1mM PMSF, 5mM beta-mercaptoethanol). The suspension was sonicated on ice at 70% amplitude with 15 sec ON and 15 sec OFF for 8 min total sonication time (Branson Sonifier). Insoluble material was removed by centrifugation for 25 min at 25,000g. In the meantime, 5 ml packed Ni-NTA resin (Qiagen) were equilibrated in lysis buffer. The beads were added to the clarified lysate and incubated under rotation for 30 min at 4°C. The bead suspension was added to a gravity flow column and the flow through was collected. Beads were washed with 40 ml lysis buffer, 120 ml wash buffer (20 mM HEPES-KOH pH8, 1 M NaCl, 0.5% NP40, 10% glycerol, 30 mM imidazole), and 80 ml lysis buffer. Proteins were eluted twice with 30ml elution buffer (20 mM TRIS-HCl pH 7.5, 300 mM NaCl, 300 mM imidazole). Two HiTrap Heparin 5ml columns (GE Healthcare) were installed in tandem on an Äkta chromatography system (GE Healthcare) and eliquibrated with 10 column volumes of HIEX buffer A (20 mM TRIS-HCl pH 7.5, 300 mM NaCl). The Ni-NTA eluate was applied and the columns and washed with 10 column volumes of HIEX buffer A. Bound proteins were eluted into fractions with a linear salt gradient of HIEX buffer A and HIEX buffer B (20 mM TRIS-HCl pH 7.5, 2 M NaCl). Fractions were analysed by SDS-PAGE and the desired ones were pooled. After concentration (Amicon Ultra-15 10 kD Spin Filters, EMD Millipore) proteins were applied to a Superdex 75 increase 10/300 GI column and eluted with the final storage buffer (HEPES-KOH pH 7.5, 250 mM KCl, 1 mM TCEP). Fractions were analysed by SDS-PAGE and the desired ones were pooled, concentrated, and stored at −80°C.

#### ORF9 purifications for phase separation assay (Figure S6C)

Recombinant ORF9 was purified using a protocol previously optimised for expression in MDG and M9ZB media (Zhou et al., 2019). Briefly, BL21-RIL DE3 *E. coli* (Agilent) were transformed with pET-6xHis-SUMO bacterial expression vectors for full-length ORF9 or C-terminally truncated ORF9 1–244 (ORF9-N). A starter culture of 30 ml grown in MDG-medium was used to inoculate 2x 1 L M9ZB-media cultures. Cultures were grown to an OD of ~2.5, chilled to 16°C, and protein expression was induced with 0.5 mM IPTG before incubation at 16°C for ~16 hours. Pelleted bacteria were lysed in lysis buffer (20 mM HEPES-KOH pH 7.5, 400 mM NaCl, 10% glycerol, 30 mM imidazole, 1 mM DTT) by sonication. Insoluble material was removed by centrifugation and 6xHis-tagged protein was purified using Ni-NTA (Qiagen) and gravity chromatography. Proteins were eluted with 30 ml elution buffer (20 mM HEPES-KOH pH7.5, 400 mM NaCl, 10% glycerol, 300 mM imidazole, 1mM DTT), dialysed against low salt buffer (150 mM NaCl) overnight in the presence of 250 μg hSENP2 protease for cleave of the 6xHis-SUMO tag. Cleaved protein was further purified by HiTrap SP ion-exchange for full length ORF9 and a combination of HiTrap Q ion-exchange and Superdex 75 size-exclusion chromatography for ORF9 1–244. Final purified fractions were pooled, concentrated and flash-frozen in liquid nitrogen for storage at −80°C.

#### cGAS

Recombinant human cGAS protein used for EMSA and cell-free cGAS activity assays was purified as described previously (Zhou et al., 2019).

### Immunoblotting

Protein concentrations in lysates were determined using Pierce BCA Protein Assay Kit (Thermo Scientific) and equalised by dilution of samples with lysis buffer. Subsequently, 4x NuPAGE LDS Sample Buffer (Invitrogen) was added and samples were incubated at 95°C for 10min. Samples were run on NuPAGE Novex 4–12% Bis-TRIS gels (Invitrogen) using NuPage MOPS-SDS running buffer (Invitrogen). Proteins were subsequently blotted onto PROTRAN Pure nitrocellulose membrane (PerkinElmer) using transfer buffer (25mM TRIS, 192mM glycine). Membranes were blocked with 5% skim milk powder (Sigma-Aldrich) in TBS containing 0.1% Tween-20 (5% milk TBS-T) for 1 hour at room temperature and were then incubated with primary antibodies in 5% milk TBS-T overnight at 4°C. Primary antibodies that bind to phosphorylated residues were diluted in 5% BSA in TBS-T. Membranes were washed thrice with TBS-T and incubated with HRP-coupled secondary antibodies in 5% milk TBS-T for 1 hour at room temperature. After three further washes with TBS-T, proteins were detected using Western Lightning Plus-ECL (PerkinElmer) and the iBright FL1000 Imaging System (Thermo Fisher) or Amersham Hyperfilm MP (GE Healthcare). If needed, antibodies were stripped from the membrane with stripping buffer (200mM glycine, 0.1% SDS, 1% Tween-20, pH 2.2) for 20min at room temperature. Membranes were washed with TBS-T, blocked as described above and re-probed.

### RT-qPCR

Cells were lysed in RLT Plus buffer and RNA was extracted using RNeasy Plus Mini Kit (Qiagen). The RNA was reverse transcribed using SuperScript IV Reverse Transcriptase (Invitrogen) and oligo-dT primers (Invitrogen). The qPCR reaction containing TaqMan Universal PCR Master Mix (Applied Biosystems) and TaqMan Primer/Probes was run on QuantStudio 7 Flex Real-Time PCR System (Thermo Fisher) with standard settings. Gene expression was analysed with the Ct method using *GAPDH* expression for normalisation. Taqman primer/probes used were: *GAPDH* (Hs02758991_g1), *IFNB1* (Hs02621180_s1), *IFI44* (Hs00951349_m1), and *IFNL1* (Hs00601677_g1). RT-qPCR for VZV transcripts was performed with SYBR GreenER master mix (Invitrogen) using primers pairs indicated in Table S3.

### Cell-free cGAS activity assay

Cell-free cGAS activity assays were performed according to a previously described procedure (Kranzusch et al., 2013). Full-length recombinant human cGAS protein (1 uM) and VACV70mer DNA (1 uM) were incubated in 20ul reaction buffer (50 mM TRIS-HCl pH 7.5, 10 mM Mg(OAc)_2_, 10 mM KCl, 1 mM DTT, 25 μM ATP, 25 μM GTP, 1 μCi α32P-ATP (PerkinElmer)) for 90 min at 37°C. Reactions were terminated for 5 min at 95°C. Leftover ATP was converted to inorganic phosphate by addition of 10 U calf-intestinal phosphatase and incubation for 20 min at 37°. 2 μl of each reaction were spotted onto a PEI Celluose F thin-layer chromatography plate (EMD Millipore) and run in 1.5 M KH_2_PO_4_. Plates were dried at room temperature and exposed to a phosphorscreen overnight. Screens were imaged using a Typhoon FLA 9500 imager (GE Healthcare). Signal intensities were quantified using Fiji 2.0.0 software (Schindelin et al., 2012) and normalized to the average signal in conditions with BSA.

### Phase separation assays

*In vitro* phase separation was performed as previously described (Du and Chen, 2018; Zhou et al., 2021). Briefly, human cGAS was labelled with AlexaFluor-488 (AF488) carboxylic acid (succinimidyl ester) (Thermo Fisher Scientific) according to manufacturer’s manuals using a molar ratio of 1:10 at 4°C for 4 h. Excess free dye was removed by dialysis against buffer (20 mM HEPES-KOH pH 7.5, 250 mM KCl, 1 mM DTT) at 4°C overnight and then human cGAS was further purified on a PD-10 desalting column (GE Healthcare) eluted with storage buffer (20 mM HEPES-KOH pH 7.5, 250 mM KCl, 1 mM TCEP) as previously described (Zhou et al., 2021).

To induce phase separation, human cGAS, ORF9, or ORF9 truncation (10 μM each) was incubated with 100 bp dsDNA (10 μM, containing 1 μM Cy3-labelled DNA) in the presence of various salt concentrations at 25°C in a total reaction volume of 20 μL. The details of proteins, nucleic acids, and salt concentrations are provided in figure legends. Reactions were placed in 384-well non-binding microplates (Greiner Bio-One) and incubated at 25°C for 30 min prior to imaging to allow condensates to settle. Fluorescence microscopy images were acquired at 25°C using a Leica TCS SP5 X (Leica Microsystems) mounted on an inverted microscope (DMI6000; Leica Microsystems) with an oil immersion 63×/numerical aperture 1.4 objective lens (HCX PL APO; Leica Microsystems). Labelled proteins were detected with excitation at 488 nm (emission at 500–530 nm) and DNA was detected with excitation at 550 nm (emission at 560–590 nm). Microscopy images were processed with FIJI (Schindelin et al., 2012), and contrast adjusted with a uniform threshold setup for each enzyme.

### Protein sequence analysis

The HSV-1 VP22 (PDB: 4XAL) and MHV68 ORF52 (PDB: 2OA5) crystal structures were used to graphically represent secondary structure features. For VZV ORF9 (UniProt accession Q4JQW6) and KSHV ORF52 (UniProt accession F5HBL8), structural features and sequence disorder was predicted by the PROFphd secondary structure prediction algorithm through submission of protein sequences to the predictprotein.org web interface (Yachdav et al., 2014).

### Data Analysis and Software

Data were analysed using Excel for Mac (Microsoft) and GraphPad Prism 8 (GraphPad Software). SnapGene (GSL Biotech) and ApE (M. Wayne Davis, The University of Utah) were utilised for DNA sequence analysis to assist cloning. Statistical analysis is detailed in the figure legends and was performed using GraphPad Prism 8. Graphs and figures were created using GraphPad Prism 8 and Adobe Illustrator CC (Adobe Systems). Immunoblot images were processed using web-based iBright Image Analysis software (Thermo Fisher). Flow cytometry data was analysed using FlowJo (FlowJo, LLC). Fiji 2.0.0 software was used to process confocal microscopy images (Schindelin et al., 2012).

