## SUPPLEMENTAL MATERIALS for "Varicella-Zoster Virus ORF9 Is an Antagonist of the DNA Sensor cGAS"

**Supplemental Figures 1-6**

**Supplemental Tables 1-3**

### Supplemental Figures

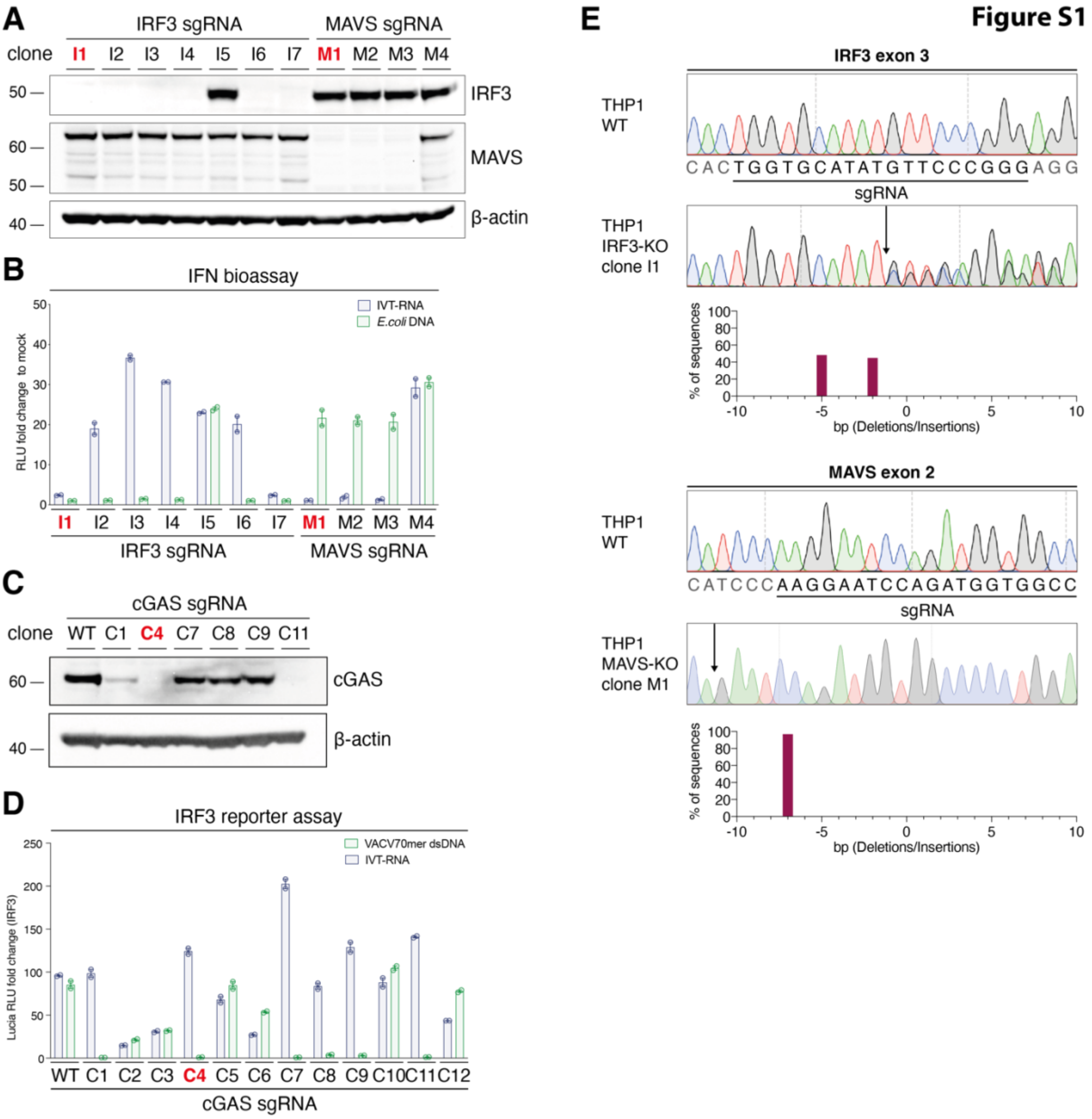


**Figure S1: IRF3, MAVS, and cGAS THP1 Dual knockout clone validation. Related to Figure 1.**
**(A, C)** THP1 Dual cells were transfected with plasmids encoding *IRF3*, *MAVS* or *cGAS* targeting sgRNAs, Cas9, and a fluorescent marker protein. Cells expressing the marker were purified by FACS and subjected to limiting dilution. Clones were expanded and tested by immunoblotting. Membranes were probed with the indicated antibodies. **(B)** Cells from the same clones as in (A) were PMA-differentiated and transfected with IVT-RNA or *E.coli* DNA. The next day, supernatants were analysed by IFN bioassay (see methods). Fold changes were calculated based on supernatants from mock-transfected cells. **(D)** Cells from clones in (C) were PMA-differentiated and transfected with IVT-RNA or VACV70mer dsDNA. The next day, luciferase activity was determined in cell supernatants. Fold changes were calculated relative to mock-transfected cells. **(E)** The *IRF3* or *MAVS* locus was PCR-amplified from genomic DNA and analysed by Sanger sequencing. Panels show electropherograms from WT cells and the knockout clones used for experiments (highlighted in red in A-B). Sequencing traces were analysed using the TIDE algorithm.
Data are from one experiment. In (B) and (D), average and range (technical duplicates) are shown.

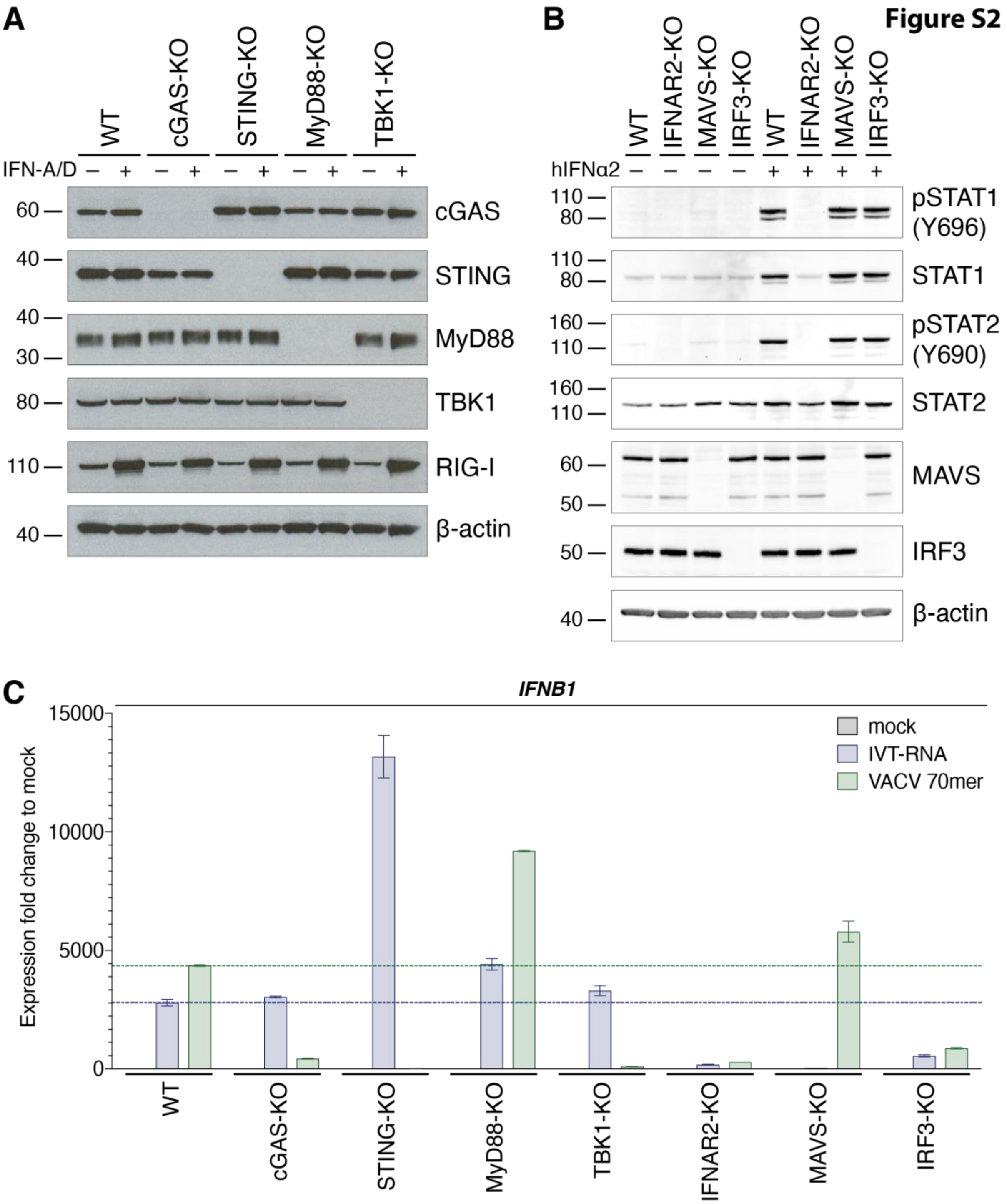


**Figure S2. Validation of THP1 knockout cells. Related to Figure 1.**
**(A,B)** The indicated THP1 cell lines were differentiated with PMA and treated or not with recombinant type I IFN as indicated. Lysates were processed for immunoblotting with the indicated antibodies. **(C)** PMA-differentiated THP1 cells were transfected with 200ng IVT-RNA or 500ng VACV 70mer DNA, or were treated with transfection reagent only (mock). RNA was extracted and RT-qPCR was performed for the *IFNB1* transcript. Horizontal lines indicate expression fold changes in WT cells.
Panel (A) is representative of three independent experiments. Panels (B) and (C) show data from one experiment. In (C), bars indicate the average and error bars the range of technical duplicates.

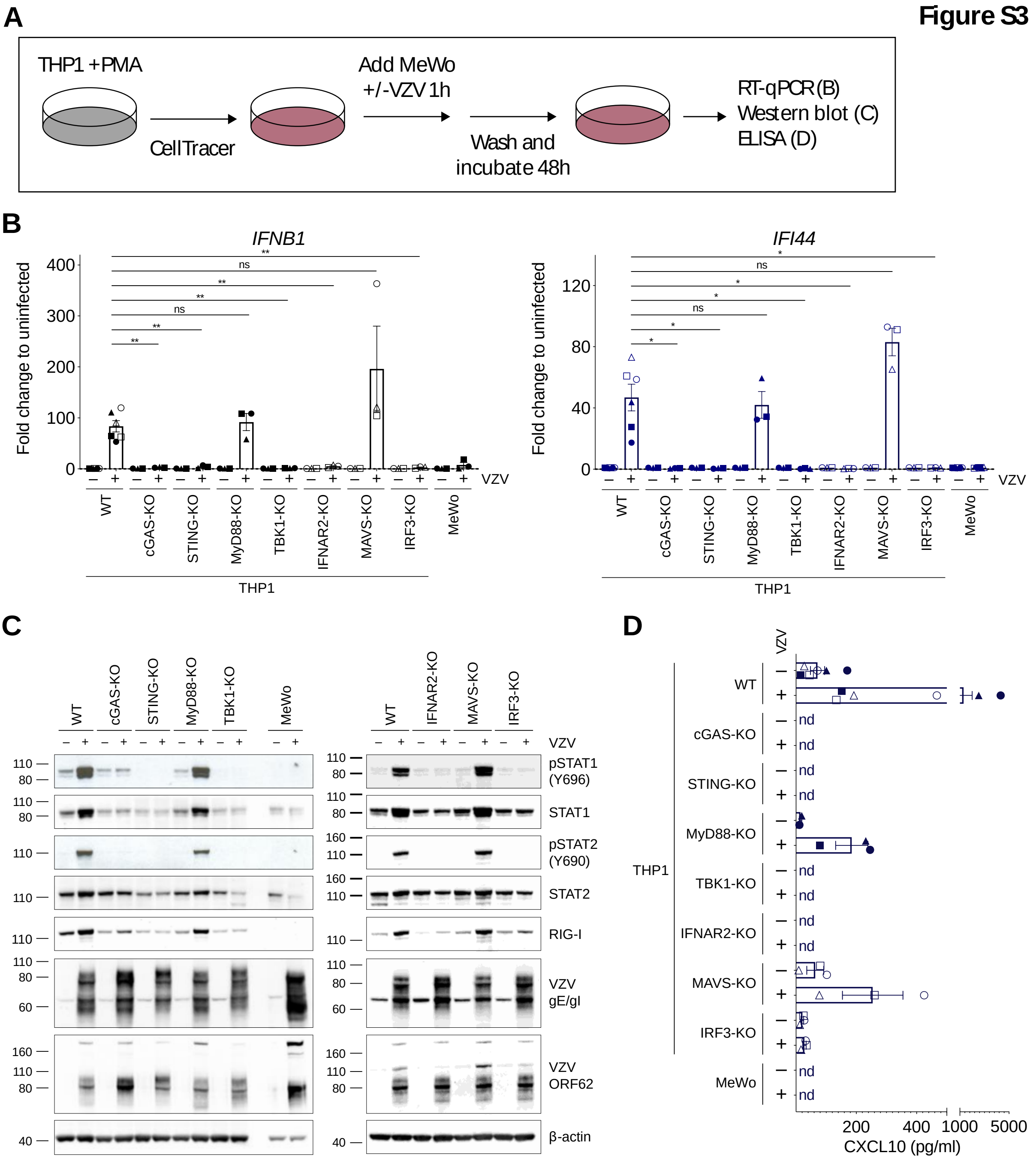


**Figure S3: The cGAS/STING DNA-sensing pathway induces type I IFNs in response to VZV infection. Related to Figure 1.**

**(A**) Schematic of the THP1-MeWo co-culture VZV infection system. See text for details. **(B)** A panel of THP1 knockout cell lines was mock infected or infected with VZV as shown in (A). The mRNA expression levels of *IFNB1* (encodes IFNβ) and *IFI44* were analysed by RT-qPCR. Expression levels were normalised to *GAPDH* and are shown as fold changes relative to levels in uninfected cells. **(C)** Cells infected as in (B) were analysed by western blot using the indicated antibodies. **(D)** Levels of CXCL10 (IP-10) in co-culture supernatants were quantified by ELISA. The different shapes of data points in (B) and (D) correspond to independent biological repeat experiments. Panels (B) and (D) show pooled data from six (THP1 WT, MeWo) or three (THP1 KOs) independent biological repeats (n=3/6 ± SEM). Panel (C) shows a representative result of three independent repeats. Statistical analysis in panel (B) was one-way ANOVA with Dunnett's multiple comparisons test. **=$p<0.01$, *=$p<0.05$, ns=not significant.

**Figure S4**

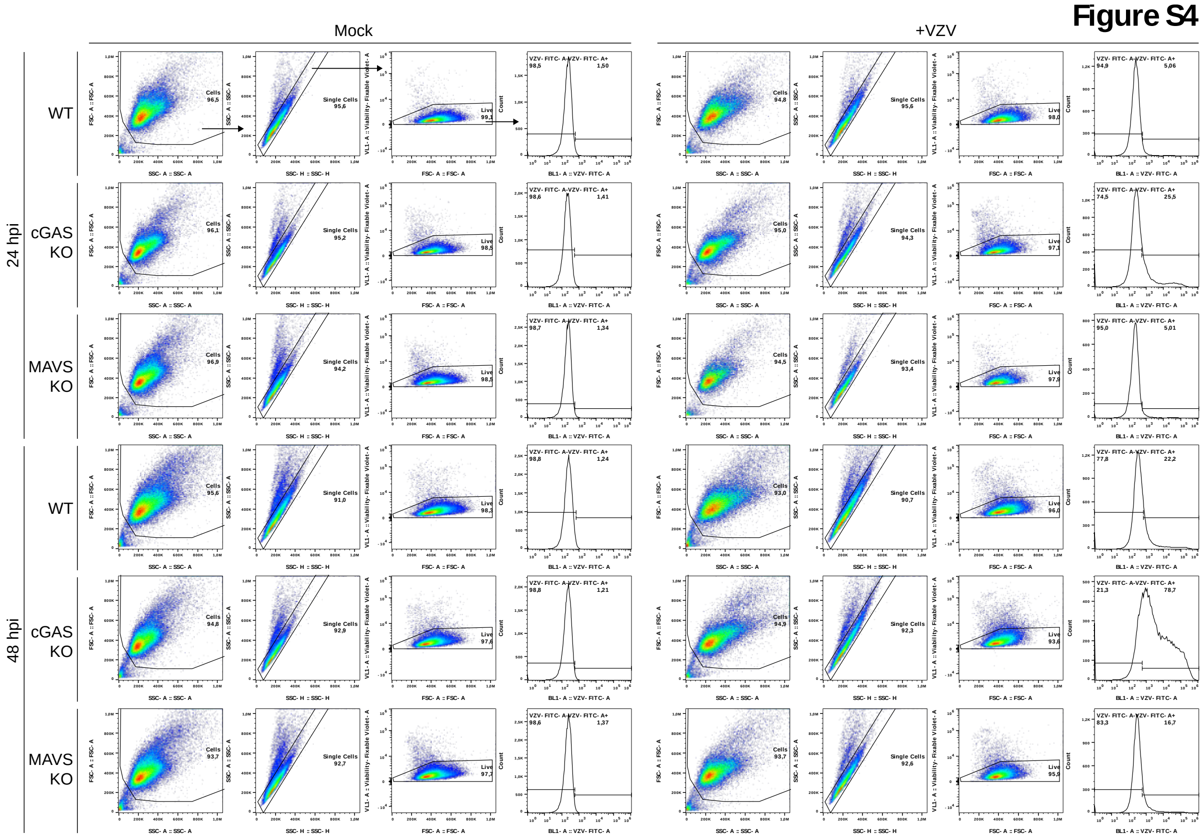


**Figure S4: Flow cytometry gating strategy for VZV-infected THP1 cells. Related to Figure 1.** Related to Figure 1C. Cells were stained with Fixable Violet Viability dye and a FITC-coupled antibody against the VZV-gE/gI heterodimer on the cell surface. FITC fluorescence was quantified after gating on single, live cells as indicated by arrows in the first row. Panel shows representative results of four independent experiments.

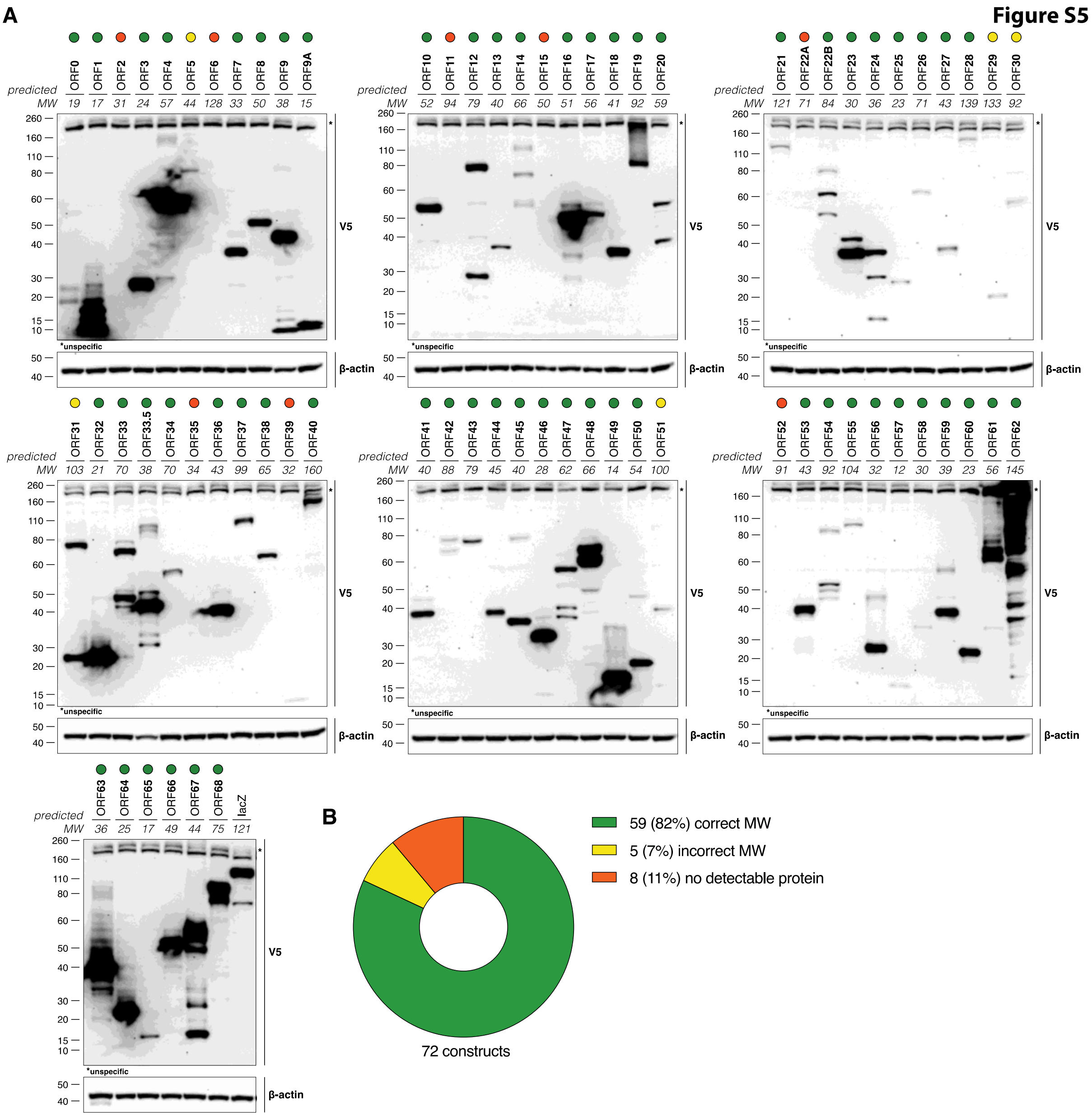


**Figure S5. VZV ORF expression library. Related to Figure 2.**
**(A)** HEK293T cells were transiently transfected with individual VZV ORF expression constructs. The next day, cell lysates were subjected to immunoblotting. Ectopically expressed proteins were detected with an antibody against the V5 tag. The predicted molecular weight (MW) is indicated in kDa. Coloured circles highlight VZV proteins expressed at the predicted MW (green), expressed at a wrong MW (orange) or not detectably expressed (red). **(B)** Summary of the data in (A).
Data in panel (A) are from one experiment.

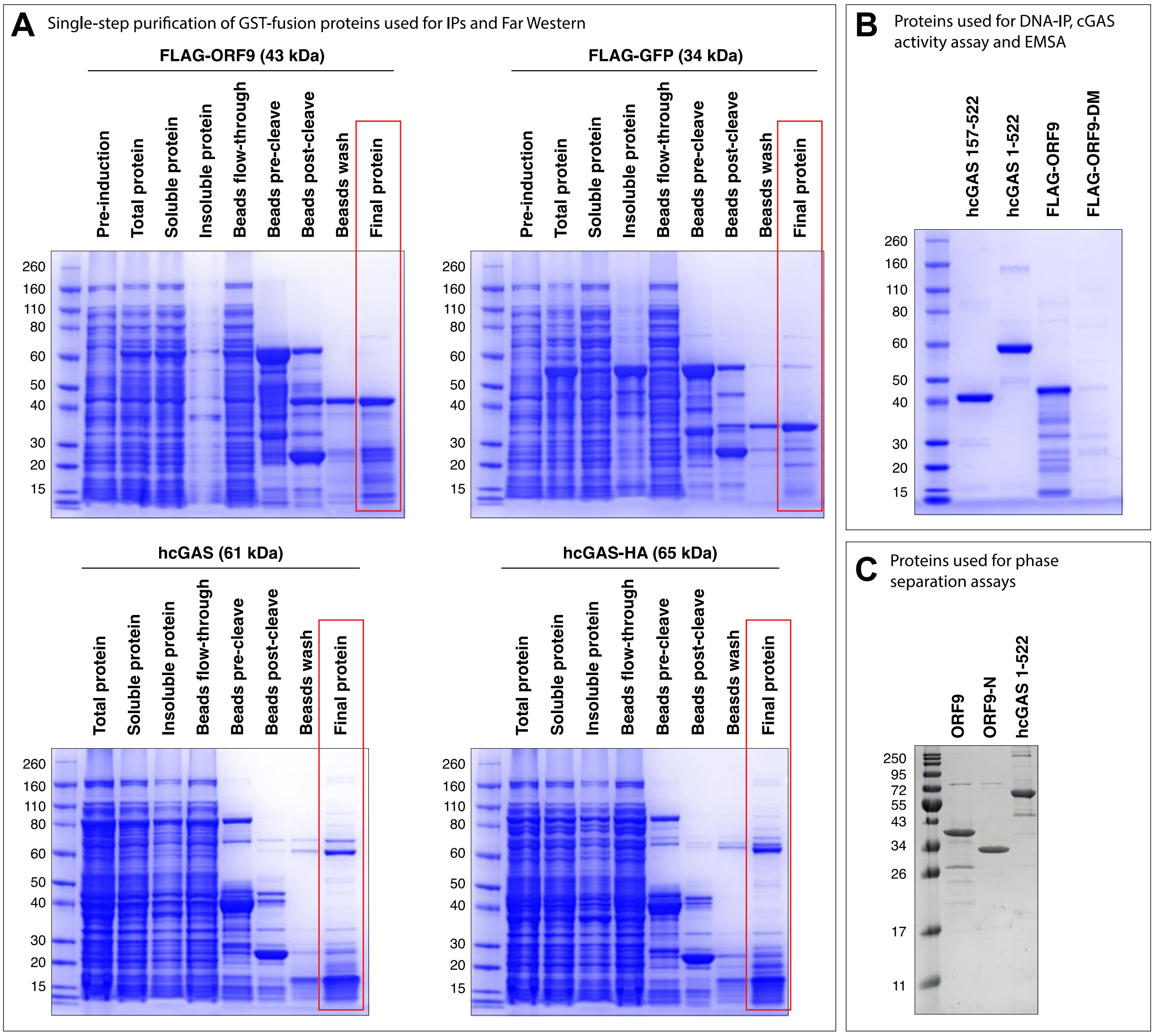


**Figure S6. SDS-PAGE analysis of recombinant protein production. Related to Figures 3, 4, 5 and 6.** **(A)** Related to Figures 3F, 3G and 4D. Images of Coomassie stained gels from production of GST-fusion recombinant proteins. See methods for details. **(B)** Related to Figures 4F, 6F and 6G. Images of Coomassie stained gels from recombinant proteins used in DNA-IPs, cGAS activity assays and agarose gel EMSA. **(C)** Related to Figure 5. Images of Coomassie stained gels from recombinant proteins used in phase separation assays. See methods for details.

### Supplemental Tables

Table S1: Primers for Cloning

| Name | F primer sequence 5'-3' | R primer sequence 5'-3' |
|---|---|---|
| hcGAS | gccgccatgcagccttggcacg | aaattcatcaaaaactggaaactcattg |
| hSTING | gccgccatgccccactccagc | agagaaatccgtgcggag |
| eGFP | gccgccatggtgagcaagggcgag | cttgtacagctcgtccatgcc |
| ORF9-STOP | gccgccatggcatcttccgacggtga | ctattttcgcgcatcagttcttgatg |
| ORF9-I-S | gccgccatggcatcttccgacggtga | ctaggcaattgcgcctgctcc |
| ORF9-II-S | gccgccatgagcgggagaccaatttccttcag | ctattttcgcgcatcagttcttgatg |
| ORF9-III-S | gccgccatgtccagcggatcggaagatg | ctattttcgcgcatcagttcttgatg |
| ORF9-IV-S | gccgccatggcatcttccgacggtga | ctataggtctgcttcattagcggcttg |
| ORF9-V-S | gccgccatgtccagcggatcggaagatg | ctataggtctgcttcattagcggcttg |
| ORF9-VI-S | gccgccatgagcgggagaccaatttccttcag | ctataggtctgcttcattagcggcttg |
| ORF9 mutant A | gcgaccgcttcacaaaagacggctgcgttatatg atggcgtaggacc | ggtcctacgccatcatataacgcagccgtcttttg tgaagcggtcgc |
| ORF9 mutant B | ctgcatggcggctacggccgcgaccgcttcacaa aag | cttttgtgaagcggtcgcggccgtagccgccatg cag |
| FLAG-ORF9-BamHI (pGEX6P1) | gggcccctgggatccgactacaaagaccatg-acggtga | gaattccggggatccctattttcgcgc -atcagttcttg |
| FLAG-GFP-BamHI (pGEX6P1) | gggcccctgggatccgactacaaagaccatg-acggtga | gaattccggggatccctacttgtacag-ctcgtccatgc |
| FLAG-ORF9/ORF9-DM (pET28a) | GGCAGCCATATGGACTACAAAGACCA TGACGGTG | GCTGCCGTCGACCTACTTGTACAGCT CGTCCATGC |

Table S2: Primers for VZV ORF PCR

| ORF | F primer sequence 5‘-3‘ | R primer sequence 5‘-3‘ |
|---|---|---|
| 0 | GCCGCCatggcgaccgtgcactactcc | tgtagttgagttgggaggttcctcgg |
| 1 | GCCGCCatgtccagggtatcggagtatgggg | ttctcgcttgcagcttgtcgc |
| 2 | GCCGCCatgcatgtaatttctgagacac | catcaatacgccctccg |
| 3 | GCCGCCatggatacaacgggagcttccg | tagtccgccgacagccg |
| 4 | GCCGCCatggcctctgcttcaattc | gcagttaaaggtactacacttaaa |
| 5 | GCCGCCatgcaggctttaggaatca | atgtttctgggagttttcac |
| 6 | GCCGCCatggataaatcctccaaacc | actcgaagttaaatttggataatt |
| 7 | GCCGCCatgcagacggtgtgtgcc | tacaagcataacatgggatttcttga |
| 8 | GCCGCCatgaacgaagcggtaattg | atgttttagtagaaaatcgacat |
| 9 | GCCGCCatggcatcttccgacggtga | ttttcgcgcatcagttcttga |
| 9A | GCCGCCatgggatcaattaccgcttcg | ccacgtgctgcgtaatacagaac |
| 10 | GCCGCCatggagtgtaatttaggaaccg | acgcgttaaaaacccaca |
| 11 | GCCGCCatgcagtcgggtcattataa | atattttcgtagtaaatgcatgg |
| 12 | GCCGCCatgttttctcggtttgcgcg | atgatgactcttaggcgtatttttcct |
| 13 | GCCGCCatgggagacttgtcatgttg | aagagccatttccatttttaggg |
| 14 | GCCGCCatgaagcggatacaaataaatttaatt | tgaacagcaacggatgca |
| 15 | GCCGCCatggccgtgaatggtgaa | cgatacatatgtaccacatagatagc |
| 16 | GCCGCCatggatttgaggtcgcgt | tttaactgtacatattacgtcagattcac |
| 17 | GCCGCCatggggctctttggactga | attccaatattttgttaatacag |
| 18 | GCCGCCatggatcagaaagattgc | taaatcgtttatcactgtgc |
| 19 | GCCGCCatggagttcaaaagaattttta | taaagcacaactggtac |
| 20 | GCCGCCatggggagtcaaccaacc | ataataacattcgttccatgtatttgt |
| 21 | GCCGCCatggaagaaccaatttgtta | agggtcactcccacttg |
| 23 | GCCGCCatgacacaacccgcatcg | caccctacgacttcttgaagc |
| 24 | GCCGCCatgtcacggagaacgtatg | tttccagaaaagcaccgc |
| 25 | GCCGCCatgtacgaatcggaaaatg | agcatccttcaatatttcatg |
| 26 | GCCGCCatggatcgggtagaatcag | gacatacttcgatagggtg |
| 27 | GCCGCCatgcatttaaagcctaccag | ccgaggaggaacaaagt |
| 28 | GCCGCCatggcgatcagaacgggg | actttgatggagaattgcttttggaa |
| 29 | GCCGCCatggaaaatactcagaagact | aatcatttccattgtaatgttcc |
| 30 | GCCGCCatggaattggatattaatcgaacattg | tgaaaacgccgggtc |
| 31 | GCCGCCatgtttgttacggcggtt | caccccgttacattctcg |
| 32 | GCCGCCatggaatcgtctaacattaacg | atcggtgtcagaatcttcat |
| 33 | GCCGCCatggctgctgaagctgac | acaccgccccaccatcat |
| 33.5 | GCCGCCatggcttctgtagcaggtaacgc | acaccgccccaccatcat |
| 34 | GCCGCCatgacggcgagatatgggtt | cggtgtggaggcaaact |
| 35 | GCCGCCatgtccgctagtcgaattcgg | cccatgggaaaacatcccgg |
| 36 | GCCGCCatgtcaacggataaaaccgat | ggaagtgttgtcctgaacg |
| 37 | GCCGCCatgtttgcgctagttttagc | tgtcagaggtattttattatattct |

| | | |
|---|---|---|
| 38 | GCCGCCatggaatttccatatcattcaac | cctttgggtttttttccc |
| 39 | GCCGCCatgaacccaccccaagcccg | aaacgaaatagatgtttttaacataacacgg |
| 40 | GCCGCCatgacaacggtttcatgt | tcgcggaagaggaaga |
| 41 | GCCGCCatggctatgccatttgagat | cacttgaatcacggcc |
| 42 | GCCGCCatgtcattgataatgtttggt | tttaataggcataaacacgg |
| 43 | GCCGCCatggaagcccatttggca | tttatggggttgggaatagag |
| 44 | GCCGCCatggaattacaacgcatatttccg | ggtggttgtaggttccggt |
| 45 | GCCGCCatgtcattgataatgtttggtcgtacg | tttaataggcataaacacggaatccg |
| 46 | GCCGCCatgtcaggccacactcca | cacatccgtgtgtggggt |
| 47 | GCCGCCatggatgctgacgacacacc | tgtcgatcctatccaatcccg |
| 48 | GCCGCCatggcacgatcgggattg | aagcaacggtttctccg |
| 49 | GCCGCCatgggacaatcttcatccag | acattttgcgcatttggaa |
| 50 | GCCGCCatgggaactcaaaagaaggg | ctcccacccactgtt |
| 51 | GCCGCCatgtctcccaacaccggg | taaactttcaaaatttaccgccccg |
| 52 | GCCGCCatggacgcaacgcagatt | taaaaacaagaagttatatgaagc |
| 53 | GCCGCCatgcagcggattcgacct | ctttacaacccgtggtgaattttt |
| 54 | GCCGCCatggccgaaataacgtct | agatcttcgatcacgtcg |
| 55 | GCCGCCatgaaaagatcaatttctgt | atacacaacgtgtacg |
| 56 | GCCGCCatgaaaaatccgcagaaattagcga | cgcgtttgcggcgtccc |
| 57 | GCCGCCatggacgtacgagaacgtaatg | acgttgaggagccttgc |
| 58 | GCCGCCatgttttcggagttgcctcc | cgttctcgtacgtccatga |
| 59 | GCCGCCatggatgtgtctggggag | tataacactccaatcgatctcg |
| 60 | GCCGCCatggcatcacataaatggttactgc | ttggcatacgcgttggaacaaa |
| 61 | GCCGCCatggataccatattagcg | ggacttcttcatcttg |
| 62/71 | GCCGCCatggatacgccgccgatgc | cccccgactctgcgggg |
| 63/70 | GCCGCCatgttttgcacctcaccggcta | cacgccatgggggggc |
| 64/69 | GCCGCCatgaatctctgcggatcccg | ggatctctcgtgggttcttgg |
| 65 | GCCGCCatggccggacaaaacacc | tccaacaaattgtgacgttat |
| 66 | GCCGCCatgaacgacgttgatgcaac | atctccaacttccattggatttg |
| 67 | GCCGCCatgtttttaatccaatgtttgat | tttaacaaacgggtttaca |
| 68 | GCCGCCatggggacagttaataaacc | ccgggtcttatctatatacaccgtgt |

Table S3: Primers for VZV RT-qPCR using SYBR Green

| | | |
|---|---|---|
| *GAPDH* | CATGGCCTTCCGTGTTCCTA | CCTGCTTCACCACCTTCTTGAT |
| *ORF40* | CCGACACGCCAGGGAACCTA | CACACCGTCAACCTGCCGTC |
| *ORF54* | TCCAACCCCTCTTCGGCTCG | GGGGATGGCCGATGGGATGT |
| *ORF63* | CCGACGCGGAATCATCGGAC | TGTTGCACCCATCCCCGTCT |
